# Activity profiling and structures of inhibitor-bound SARS-CoV-2-PLpro protease provides a framework for anti-COVID-19 drug design

**DOI:** 10.1101/2020.04.29.068890

**Authors:** Wioletta Rut, Zongyang Lv, Mikolaj Zmudzinski, Stephanie Patchett, Digant Nayak, Scott J. Snipas, Farid El Oualid, Tony T. Huang, Miklos Bekes, Marcin Drag, Shaun K. Olsen

## Abstract

In December 2019, the first cases of a novel coronavirus infection causing COVID-19 were diagnosed in Wuhan, China. Viral Papain-Like cysteine protease (PLpro, NSP3) is essential for SARS-CoV-2 replication and represents a promising target for the development of antiviral drugs. Here, we used a combinatorial substrate library containing natural and a wide variety of nonproteinogenic amino acids and performed comprehensive activity profiling of SARS-CoV-2-PLpro. On the scaffold of best hits from positional scanning we designed optimal fluorogenic substrates and irreversible inhibitors with a high degree of selectivity for SARS PLpro variants versus other proteases. We determined crystal structures of two of these inhibitors (VIR250 and VIR251) in complex with SARS-CoV-2-PLpro which reveals their inhibitory mechanisms and provides a structural basis for the observed substrate specificity profiles. Lastly, we demonstrate that SARS-CoV-2-PLpro harbors deISGylating activities similar to SARS-CoV-1-PLpro but its ability to hydrolyze K48-linked Ub chains is diminished, which our sequence and structure analysis provides a basis for. Altogether this work has revealed the molecular rules governing PLpro substrate specificity and provides a framework for development of inhibitors with potential therapeutic value or drug repositioning.

## Introduction

The global epidemic of three coronaviruses have emerged in this century so far. In November 2002 in Foshan, China, the first known case of human infected with severe acute respiratory syndrome coronavirus (SARS-CoV) has been reported (*1*). By July 2003, more than 8,000 SARS cases were detected in 27 countries. The main symptoms of SARS-CoV infection were influenza-like and included fever, headache, malaise, shivering and diarrhea. Only a few cases of infection occurred between December 2003 and January 2004 (*2*). The implementation of measures infection control has ended the global SARS outbreak. Ten years after the SARS pandemic a new coronavirus – Middle East respiratory syndrome coronavirus (MERS-CoV) was diagnosed in Saudi Arabia man (*3*). Due to international travels of infected people, MERS-CoV has spread worldwide. 2502 laboratory-confirmed cases of MERS-CoV infection were reported from September 2012 to the end of December 2019, including 858 associated deaths. In December 2019 a novel coronavirus - severe acute respiratory syndrome coronavirus 2 (SARS-CoV-2) formerly known as the 2019 novel coronavirus (2019-nCoV) was identified in Wuhan, China (*4*, *5*). Current studies indicate that this coronavirus is similar to SARS-CoV. Although these three coronaviruses - SARS-CoV, MERS-CoV, and SARS-CoV-2 are identified as a highly pathogenic into the human population, there is no effective antiviral treatment. Therefore, current studies are focused on rapid development of vaccines and antiviral drugs to prevent and treat coronavirus infection.

One of the attractive antiviral drug targets is the SARS-CoV encoded cysteine protease – papain-like protease (PLpro) (*6*). This enzyme recognizes the tetrapeptide LXGG motif found in-between viral proteins nsp1 and nsp2, nsp2 and nsp3, and nsp3 and nsp4 (nsp1/2, nsp2/3, nsp3/4) (*7*, *8*). The hydrolysis of the peptide bond on the carboxyl side of glycine at the P1 position leads to the release of nsp1, nsp2, and nsp3 proteins, which are essential for viral replication. The *in vitro* studies have shown that SARS-CoV-PLpro harbors two other proteolytic activities, removal of ubiquitin (Ub) and ubiquitin-like protein ISG15 (interferon-induced gene 15) from cellular proteins (*9*–*11*). Ubiquitinated and ISGylated substrates are more efficiently hydrolyzed by SARS-CoV-PLpro than small substrates containing C-terminal LRGG motif (*11*, *12*). These results indicated a more complex mechanism of substrate recognition than only the interaction of S4-S1 pockets of enzyme with tetrapeptide fragment. Further studies revealed that SARS-CoV-PLpro possess two distinct Ub binding subsites (SUb1 and SUb2) and recognize Lys48-linked polyUb chains for polyubiquitin chain editing and/or deubiquitination of polyubiquitinated proteins (*13*–*15*).

Due to the deubiquitinating and deISGylating activities of SARS-CoV PLpro, this enzyme performs significant role in the innate immune response during viral infection (*16*, *17*). SARS-CoV-PLpro is involved in inhibiting the production of cytokines and chemokines, that are responsible for the activation of the host innate immune response against viral infection (*18*–*20*). For these reasons, this enzyme is an important molecular target in the design of SARS-CoV antiviral drugs. Despite substantial research efforts in the development of SARS-CoV inhibitors, efficacy data of these compounds from clinical trials are missing (*21*–*23*). Nevertheless, we hypothesize that information gained over past years for the SARS-CoV-PLpro could be immediately translated into the timely study of SARS-CoV-2-PLpro to accelerate new antivirals development and drug retargeting approaches. The optimal method to check level of similarity in binding site architecture between two enzymes is positional scanning technology. We have recently developed a novel chemical approach, called HyCoSuL (Hybrid Combinatorial Substrate Library), to dissect a broad substrate specificity of proteolytic enzymes (*24*). Application of this approach SARS-CoV-2-PLpro facilitated the development of irreversible inhibitors (VIR250 and VIR251) harboring a high degree of selectivity for SARS PLpro variants versus other proteases. Crystal structures of VIR250 and VIR251 in complex with SARS-CoV-2-PLpro reveal their inhibitory mechanisms and provide a structural basis for the observed substrate specificity profiles. We also examined processing of Ub and Ubl by SARS-CoV-2 PLpro and found that it harbors deISGylating activities similar to SARS-CoV-1-PLpro but its ability to hydrolyze K48-linked Ub chains is significantly diminished. Altogether this work has revealed the molecular rules governing PLpro substrate specificity, provides a framework for development of inhibitors with potential therapeutic value or drug repositioning, and reveals intriguing differences in the biochemical functions of SARS-CoV-1 and 2 PLpro that may contribute to differences in their biology.

## Results

### Substrate specificity profile

SARS-CoV-2 PLpro recognizes the tetrapeptide LXGG motif found in-between viral proteins nsp1 and nsp2, nsp2 and nsp3, and nsp3 and nsp4 (Figure 1A) (*7*, *8*). Hydrolysis of the peptide bond on the carboxyl side of glycine at the P1 position leads to the release of nsp1, nsp2, and nsp3 proteins, which are essential for viral replication. SARS-CoV-2 PLpro also harbors deubiquitinating and deISGylating activities and recognizes the conserved LRGG motif at the C-terminus of these proteins. (Figure 1A). Our previous studies of SARS-CoV-1 PLpro substrate preferences using a combinatorial substrate library containing only natural amino acids revealed that this protease recognizes LXGG motif at P4-P1 positions with broad substrate specificity at P3 position (*25*). These results suggest that more detailed mapping of binding pocket architecture should facilitate design of new, active substrates as well as optimal peptide sequences for inhibitor development efforts. To achieve this goal we developed a defined and combinatorial substrate library (HyCoSuL) containing wide variety of nonproteinogenic amino acids (*24*). Since tetrapeptide fluorogenic substrates are not very efficiently hydrolyzed by enzymes exhibiting deubiquitinating activity, we designed and synthesized the P2 defined library with a general structure of Ac-LRXG-ACC (X – 19 natural and 109 unnatural amino acids) and a hybrid combinatorial substrate library, where three positions were fixed and one position contains an equimolar mixture of 19 amino acids (Mix), (P3 sublibrary: Ac-Mix-P3-Gly-Gly-ACC, P4 sublibrary: Ac-P4-Mix-Gly-Gly-ACC; P3 and P4 – a natural or unnatural amino acid) (*26*). By design of libraries with tailored peptide scaffold toward DUBs we could reach the highest possible concentration of individual fluorogenic substrates in each sublibrary during the assay.

**Fig. 1.**
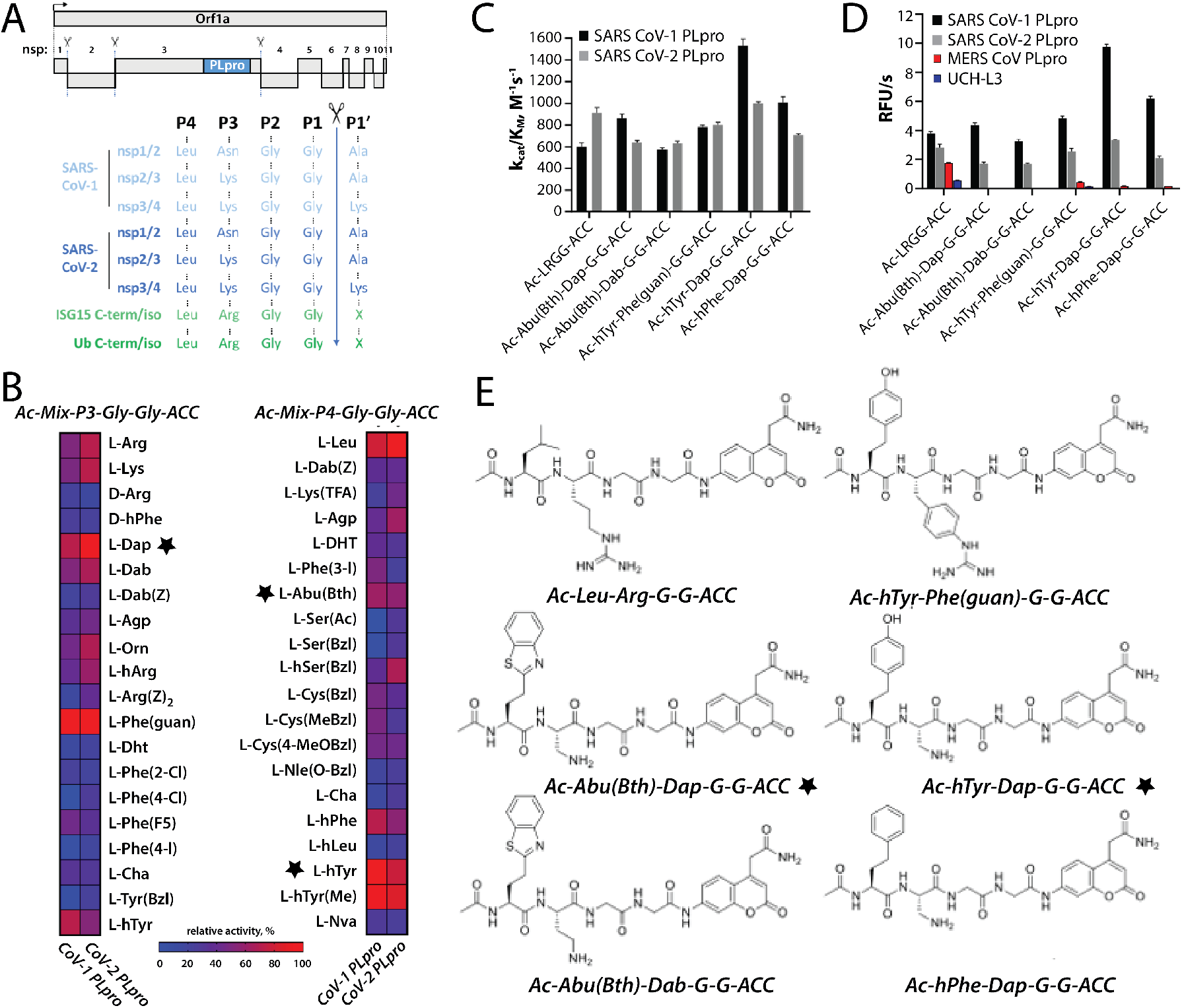
Activity profiling of SARS-CoV-1 and 2 PLpro protease. **(A)** top, Schematic representation of Orf1a from SARS-hCoV-2, indicating the processed non-structural proteins (nsps). PLpro cleavage sites are indicated by scissors. Bottom, sequence of alignment of P4-P1’ aminoacids of the indicated PLpro cleavage sites. **(B)** SARS-CoV-1 PLpro and SARS-CoV 2-PLpro substrate specificity profiles at the P3 and P4 positions presented as heat maps. Asterisk indicates P3 and P4 groups used throughout the study. **(C)** k_cat_/K_M_ for tetrapeptide fluorogenic substrates toward SARS-CoV-1 PLpro and SARS-CoV-2 PLpro. **(D)** The rate of tetrapeptide substrate hydrolysis by DUBs. ([S]=10 μM; SARS-CoV PLpro concentration, 0.2 μM; SARS-CoV-2 PLpro concentrations, 0.1 μM; MERS-CoV PLpro concentration, 2.5 μM; UCH-L3 concentration, 8.8 μM). **(E)** Tetrapeptide substrate structures.

P2 library screening revealed that SARS-CoV and SARS-CoV-2-PLpro possess very high substrate specificity at this position – only glycine can be accepted (Supplementary Figure 1). Both proteases exhibit a broad substrate preference at P3 position (Figure 1B). The S3 pocket of SARS-CoV and SARS-CoV-2-PLpro can tolerate not only positively charged residues like Phe(guan), Dap, Dab, Arg, Lys, Orn, and hArg, but also hydrophobic amino acids, such as hTyr, Phe(F5), Cha, Met, Met(O), Met(O)_2_, D-hPhe (amino acid structures presented in Table S1 supplemental information). These enzymes do not recognize acidic residues and most D-amino acids (the exception are D-Arg, D-hPhe, D-Lys, and D-Phg). The S4 pocket of SARS-CoV and SARS-CoV-2-PLpro can accommodate hydrophobic residues only, among natural amino acids, practically only leucine can be tolerated (being the best hit for SARS-CoV-2-PLpro) (Figure 1B). SARS-CoV-PLpro recognized two unnatural residues better than leucine at P4 position (hTyr and hTyr(Me)). Other bulky amino acids are also accepted (≥ 30%, hPhe, Abu(Bth), Phe(3-I), Cys(Bzl), Cys(MeBzl), Cys(4-MeOBzl), hSer(Bzl), and Dht) (Figure 1B).

### Design and kinetic analysis of tetrapeptide fluorogenic substrates

To validate the library-screening data we designed optimal tetrapeptide fluorogenic substrates to find optimal sequences recognized by SARS-CoV and SARS-CoV-2-PLpro. We analyzed both SARS PLpro substrate specificity profile at the P4-P2 positions and selected the most preferred amino acids (P2: Gly; P3: Dap, Phe(guan); P4: hTyr, hPhe, and Abu(Bth)) (Figure 1E). Kinetic analysis revealed that some designed substrates were better recognized by SARS-CoV-1 PLpro with Ac-hTyr-Dap-Gly-Gly-ACC being almost 2.5 times more efficiently cleaved than endogenous Ac-LRGG-ACC. In case of SARS-CoV-2-PLpro, we did not find significant difference between Ac-LRGG-ACC and all tested substrates (Figure 1C). It is important to notice that substitution of Arg in P3 position by relatively small Dap did not affect binding to S3 pocket and yields very good substrates (Figure 1C). Thus, data obtained from combinatorial screening very well translate into individual substrates and demonstrate very high level of similarity between two investigated enzymes.

Next, we wanted to see if incorporation of unnatural amino acids in P4 and P3 positions of peptide sequence can result in selective tetrapeptide substrates. We tested the substrates with four enzymes that exhibit deubiquitinating activity – SARS-CoV-PLpro, SARS-CoV-2-PLpro, MERS-CoV-PLpro and human DUB UCH-L3. We have found that none of the substrates with unnatural amino acids in the sequence were significantly recognized at 10μM either by MERS-CoV-2 PLpro (2.5 μM) nor human DUB UCH-L3 (8 μM) (Figure 1D). In line with previous data, Ac-LRGG-ACC was recognized by all four enzymes (Figure 1D).

### Development of PLpro inhibitors

To further analyze selectivity of peptide sequences with unnatural amino acids we converted two substrates (Ac-hTyr-Dap-Gly-Gly-ACC and Ac-Abu(Bth)-Dap-Gly-Gly-ACC) into inhibitors by exchanging the fluorescent tag to a reactive group – vinylmethyl ester (VME). VME group was selected due to its broad reactivity toward DUBs (inhibitor selectivity is determined by tetrapeptide sequence). The results from kinetic analysis of SARS-CoV-PLpro and SARS-CoV-2-PLpro inhibitors reflected those of substrate hydrolysis (Figure 2A, B). Ac-hTyr-Dap-Gly-Gly-VME (hereafter referred to as VIR251) was more potent but less selective inhibitor toward these enzymes than Ac-Abu(Bth)-Dap-Gly-Gly-VME (hereafter referred to as VIR250). Importantly, both compounds exhibit high selectivity for SARS-PLpro variants and robustly inhibit both SARS-CoV-PLpro, SARS-CoV-2-PLpro activities. In contrast, practically no inhibition of human UCH-L3 and only a slight inhibition of MERS-PLpro was observed (Figure 2A, B). Further, incubation of HeLa lysates with Ub-VME yields a crosslinking profile that is unaltered by titrations of VIR250 or VIR251 (Figure 2C). Since a major cross-linking target of Ub-VME is known to be human DUB enzymes, this data suggests that VIR250 and VIR251 do not cross-react with human DUBs. This is an important finding in search for a selective antiviral molecule with minimal cross-reactivity with human DUBs.

**Fig. 2.**
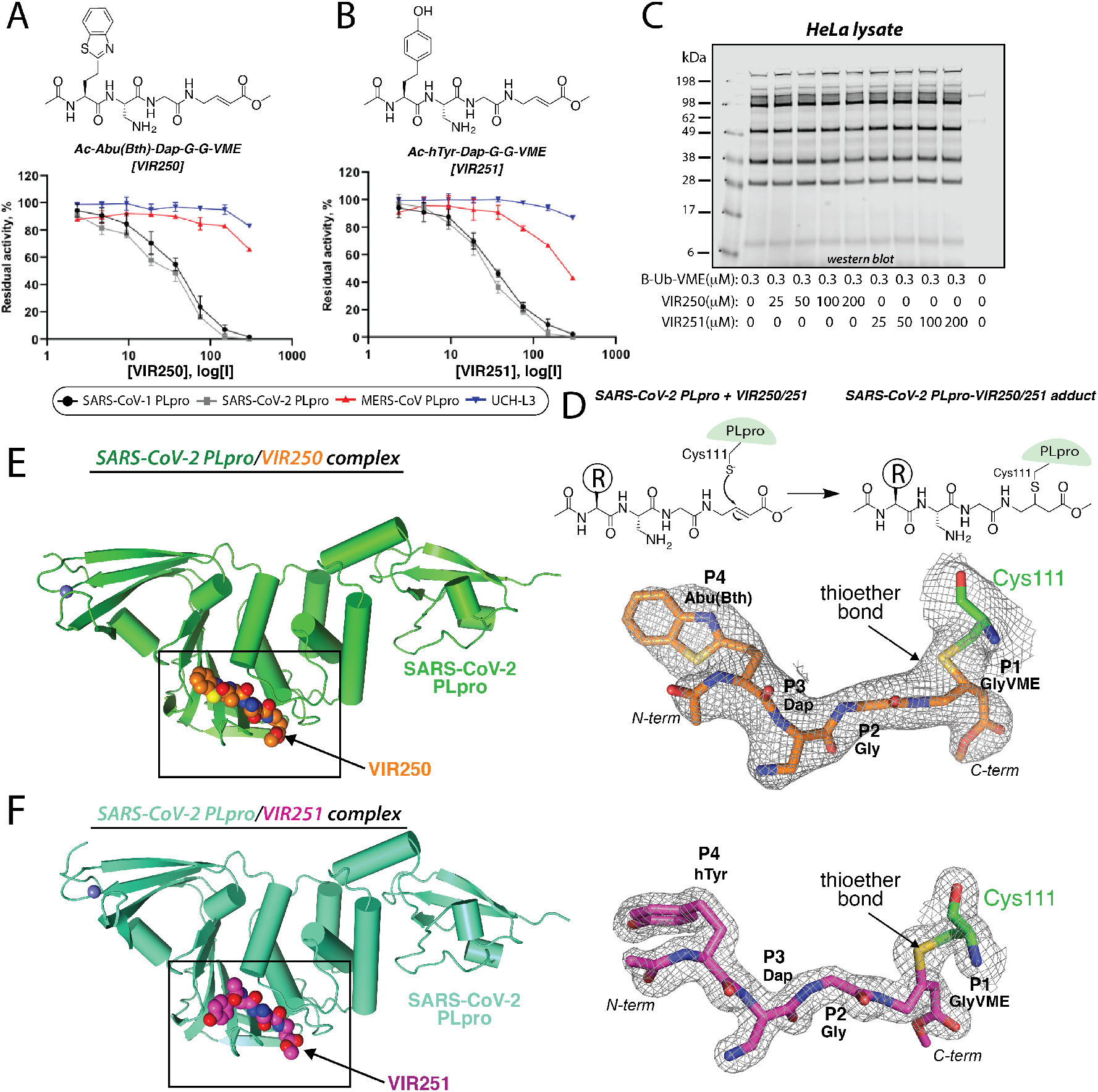
Characterization of non-natural amino acid-containing inhibitors VIR250 and VIR251 and their crystal structures in complex with SARS-CoV-2 PLpro. **(A, B)** DUBs inhibition by designed inhibitors ([I]=2.3-300 μM; SARS-CoV-PLpro concentration, 0.3 μM; SARS-CoV-2-PLpro concentration, 0.1 μM; MERS-CoV PLpro concentration, 2.5 μM; UCH-L3 concentration, 8 μM). **(C)** HeLa lysate selectivity assay. HeLa cell lysate first incubated for 30 minutes with the indicated inhibitor and next for 30 minutes with Biotin-Ub-VME followed by SDS-page and western blot using streptavidin Alexa Fluor 647. **(D)** Schematic of the crosslinking of VIR250 and VIR251 to SARS-CoV-2 PLpro. **(E)** *left*, SARS-CoV-2 PLpro is shown as green ribbon representation and VIR250 is shown as spheres with carbon (orange) nitrogen (blue) oxygen (red) and sulfur (yellow). *right* 2F_o_F_c_ electron density map (1.0 σ) for VIR250 is shown as gray mesh. Carbon atoms of catalytic Cys111 of SARS-CoV-2 PLpro are shown as green sticks. The thioether linkage between Cys111 and VIR250 is indicated with an arrow. **(F)** SARS-CoV-2 Plpro/− VIR251 structure presented as in E except with SARS-CoV-2 PLpro and VIR251 carbon atoms colored cyan and magenta, respectively. The electron density map is contoured at 1.5 σ.

### Structures of CoV-2-PLpro in complex with VIR250 and VIR251

We next set out to determine crystal structures of SARS-CoV-2-PLpro in complex with VIR250 and VIR251 in order to gain insights into the molecular mechanism by which these molecules inhibit SARS-CoV-2-PLpro activity as well as the basis for the observed substrate selectivity profile. Catalytic cysteine 111 of CoV-2-PLpro engages in Michael Addition to the β-carbon of the vinyl group of the VME warheads of VIR250 and VIR251, resulting in formation of a covalent thioether linkage (Figure 2D). Large-scale cross-linking reactions yielded CoV-2-PLpro-VIR250 and CoV-2 PLpro-VIR251 complexes of yield and purity sufficient for growth of diffraction quality crystals. The structure of CoV-2-PLpro in complex with VIR250 (Figure 2E) was determined by molecular replacement using the recently determined structure of apo CoV-2 PLpro (PDB: 6W9C) and was resolved to 2.79 Å resolution with R/R_free_ values of 0.230/0.195 (Table S2). This structure was used as the molecular replacement search model for determination of the structure of CoV-2-PLpro in complex with VIR251 (Figure 2F). The CoV-2 PLpro/VIR251 structure was resolved to 1.65 Å resolution and refined to R/R_free_ values of 0.196/0.170 (Table S2).

Comparison of apo CoV-2-PLpro to CoV-2-PLpro/VIR250 and CoV-2-PLpro/VIR251 complexes reveal similar overall structures with the exception of the β14-β15 loop which is situated proximal to the active site and undergoes a conformational change in that is likely due to inhibitor binding (Supplementary Figure 2) (see below). This analysis shows that there are also slight rigid body rotations of the finger and ubiquitin-like domains of CoV-2-PLpro which are likely due to crystal packing effects. Analysis of the structures reveal extensive electron density projecting from the caytalytic Cys111 side chain of CoV-2-PLpro into which all the atoms of VIR250 and VIR251 could unambiguously be placed (Figures 2E, F). Further, the covalent bond between Cys111 and both VIR250 and VIR251 are clear (Figures 2E, F). As anticipated, both VIR250 and VIR251 inhibitors occupy the S4-S1 pockets of CoV-2-PLpro in proximity to the active site and adopt similar structures with the exception of the orientation of the P4 substituents which will be discussed in greater detail below. The P4 position is the only region of chemical divergence between VIR250 and VIR251, with an Abu(Bth) in VIR250 and an h-Tyr in VIR251 (Figures 2A, B).

### Molecular recognition of VIR250 and VIR251

Analysis of the CoV-2-PLpro/VIR250 and CoV-2-PLpro/VIR251 complexes reveals a similar network of interacting residues with ~560 Å^2^ from a total of ~775 Å^2^ solvent accessible area of VIR251 and ~600 Å^2^ from a total of ~800 Å^2^ solvent accessible area of VIR250 buried upon complex formation. With the exception of the P4 positions of VIR250 and VIR251, which engage largely in hydrophobic interactions with CoV-2-PLpro, the majority of interactions at the P1-P3 positions of both inhibitors are mediated through polar interactions and hydrogen bonds (Figures 3A, B). At the P1 position of VIR250, GlyVME is covalently linked via thioether bond to catalytic Cys111 of CoV-2-PLpro and engages in a backbone-backbone hydrogen bond to Gly271 (Figure 3A). At the P2 VIR250 position, Gly engages in two backbone-backbone hydrogen bonds to Gly163 and van der Waals contacts to Leu163 and Tyr164 of CoV-2-PLpro and P3 Dap of VIR250 participates in a backbone-backbone hydrogen bond with Gly271 (Figure 3A). The network of backbone-backbone hydrogen bonds participated in at the P3-P1 positions of VIR250 are fully conserved in VIR251 (Figure 3B). In contrast, while the methylester group from the GlyVME warhead of VIR250 engages in a hydrogen bond with His272 from the catalytic triad of CoV-2-PLpro, the corresponding methylester of VIR251 participates in hydrogen bonds with Trp106 and Asn109 side chains, which are proposed to contribute to oxyanion hole stabilization (Figures 3A, B). Trp106 adopts a different conformation and is poorly ordered in the VIR250 complex (Figure 3A).

**Fig. 3.**
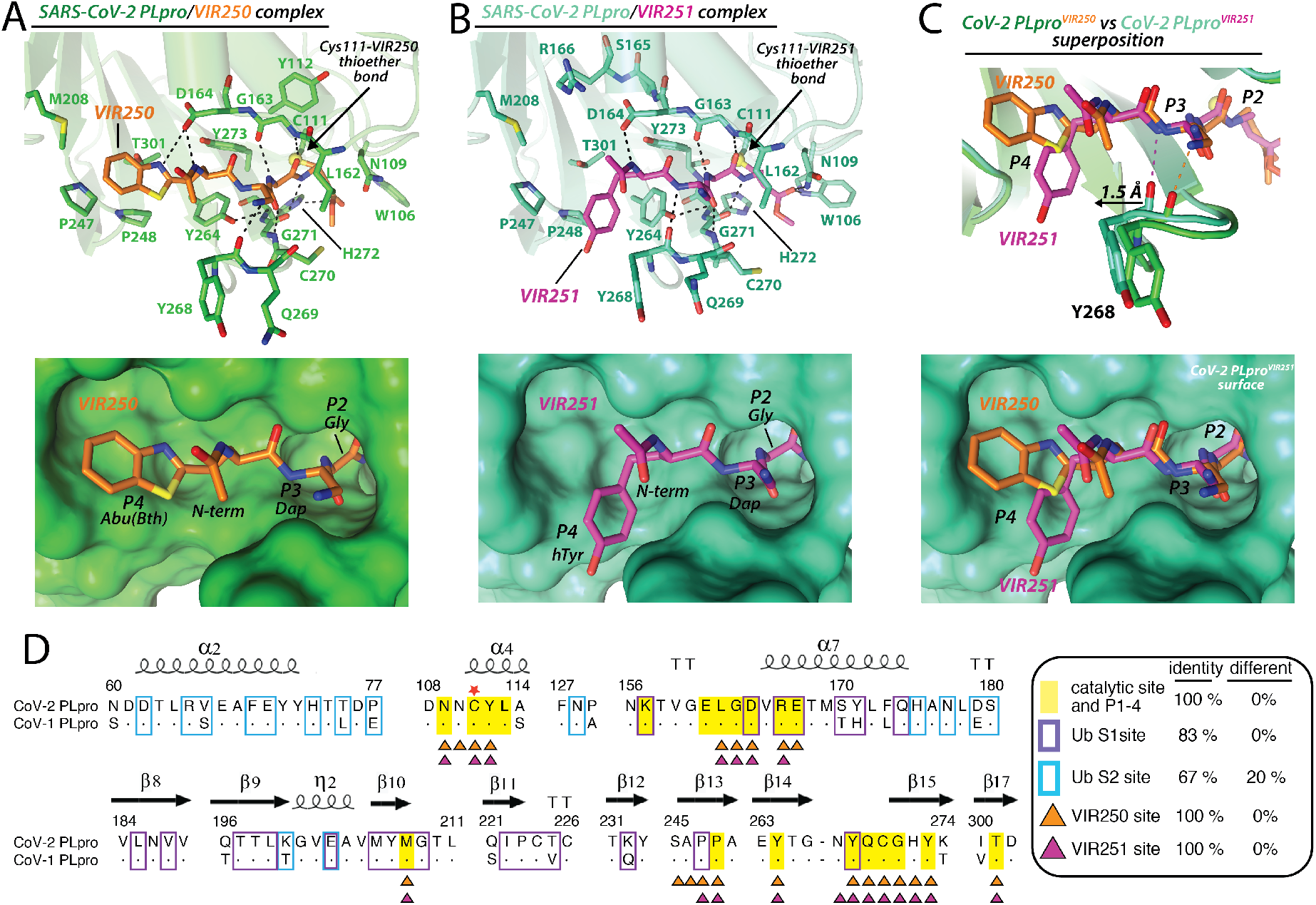
Molecular recognition of VIR250 and VIR251 by SARS-CoV-2 PLpro. **(A)** *top*, Network of contacts between SARS-CoV-2 PLpro (green) and VIR250 (orange) with involved residues shown as stickes with red oxygen atoms, blue nitrogen atoms, and yellow sulfur atoms. Hydrogen bonds are indicated by dashed lines. *bottom*, The VIR250 binding pocket of CoV-2 PLpro is shown as green surface with VIR250 shown as stricks. The P2-P4 positions are labeled. **(B)** Network of contacts between SARS-CoV-2 PLpro (cyan) and VIR250 (magenta) shown as in A. **(C)** Superimposition of the CoV-2 PLpro/VIR250 and CoV-2 PLpro/VIR251 structures presented as in A and B. Only the surface of CoV-2 PLpro from the PLpro/VIR251 structure is shown in the bottom panel. **(D)** Structure-based sequence alignment CoV-1 and CoV-2 PLpro with secondary structure was shown above sequence. Conserved residues (dots), catalytic cysteine (red star). Residues buried at the CoV-2 PLpro/VIR250 and CoV-2 PLpro/VIR251 interfaces are indicated by orange and magenta triangles below the alignment, respectively. Residues involved in contacts with the S1 Ubiquitin residue 73-76 (P1-P4), S1 Ub, and S2 Ub based on the SARS-CoV-2 PLpro/K48 diUbiquitin structure (PDB: 5E6J) are shaded yellow and boxed purple and cyan, respectively.

Intriguingly, there are significant differences in how the side chains of the P3 and P4 positions of VIR250 and VIR251 engage CoV-2-PLpro. The side chain amine of Dap at the P3 position of VIR250 engages in a hydrogen bond with the backbone carbonyl oxygen of Tyr268 and the P4 Abu(Bth) projects toward Met208, Pro247, Pro248, and Thr301 where it engages in a network of van der Waals interactions (Figure 3A). In contrast, it is the backbone amine of P3 Dap that engages in the hydrogen bond to the carbonyl oxygen of Tyr268, and surprisingly, hTyr at the P4 position projects towards the opposite side of the S4 pocket compared to Abu(Bth) from VIR250 by extending towards Pro248, Tyr264, and Tyr268 of CoV-2-PLpro and participating in a distinct network of van der Waals interactions (Figure 3B). This new network of interactions is facilitated by a 1.5 Å shift of the β14-β15 loop (Asn267, Tyr268, and Gln269) towards the hTyr of VIR251 (Figure 3C) thereby facilitating many novel contacts which would be unable to occur in the absence of this shift. Notably, all of the CoV-2-PLpro residues involved in contacts to both VIR250 and VIR251 are fully conserved in SARS CoV-1-PLpro, and the overall structures of the two SARS PLpro variants are very similar in the catalytic site of the enzyme which likely accounts for the ability of these inhibitors to target both enzymes (Figure 3C, 4G).

In terms of how our structures correlate with the observed substrate selectivity profiles described above, P2 dependence on Gly is the result of residues from the β14-β15 and α5-α6 loops of CoV-2-PLpro (notably Leu162, Tyr 264, Cys270, Gly271, and Tyr273) clamping down on top of the P2 position leaving no room for side chain atoms at the R position (Figures 3A-C). The preference for positive and hydrophobic residues and selection against acidic residues at the P3 position is likely the result of its broader pocket and proximity to the acidic carbonyl oxygens of Tyr268, Gln269, and Leu162, the side chain of Asp164, as well as the hydrophobic side chains of Leu162 and Tyr268 (Figures 3A-C). At the P4 position the strong preference for bulky hydrophobic residues can be explained by the hydrophobic nature of the P4 binding pocket that is largely formed by residues Met208 Pro247, Pro248, Tyr264, and Tyr268 (Figures 3A, B). Notably, the very deep and broad nature of the S4 pocket of SARS-CoV-2-PLpro has been exploited by the Abu(Bth) and h-Tyr sidechains at P4 of VIR250 and VIR251, which as noted above project towards different ends of the S4 pocket and engage in distinct networks of contacts (Figure 3C). With that said there remain regions at the deepest parts of this pocket, particularly an acidic patch formed by Asp164, Tyr273, and Thr301 that could potentially be exploited for development of more potent inhibitors.

### Processing of Ub and Ubl variants by CoV-1 and CoV-2-PLpro

Studies carried out for SARS-CoV-1 PLpro revealed that this enzyme has ubiquitin binding domain and efficiently process full ubiquitin fluorogenic substrates (*12*). We wanted to see if this is also the case for SARS-CoV-2-PLpro. To that end, we used ubiquitin activity-based probe (ABP) for labeling of both enzymes. In this ABP, biotin was used as detection tag and VME as an irreversible warhead that cross-links to the catalytic cysteines. To test its sensitivity, we performed SDS-PAGE analysis followed by nitrocellulose membrane transfer and visualization with fluorescent streptavidin (Figure 4A). We observed significant labeling of both proteases by Biotin-Ub-VME at a concentration twice higher than the enzyme concentration (200 nM), however SARS-CoV-1 PLpro was more efficiently labeled compared to SARS-CoV-2 PLpro. We next decided to explore the Ub and Ub-like protein (Ubl) substrate specificity of SARS-CoV-2 PLpro by testing its reactivity with a panel of Ub/Ubl ABPs. This panel of probes includes K48-linked diUb and K63-linked diUb with propargylamide (PA) warheads at their distal ends, mono Ub-PA, and ISG15 C-terminal domain (CTD)-vinyl pentynyl sulfone (VPS). The results of this analysis indicated a preference of SARS-CoV-2 PLpro for recognition of ISG15 and monoUb and a poor ability to recognize K48-linked and K63-linked diUb (Figure 4B). As expected, preincubation of SARS-CoV-2 PLpro with VIR251 completely blocked its ability to react with all of the probes (Figure 4B). The apparent lack of reactivity with the K48-linked diUb-PA probe and the increased processing of ISG15^CTD^-VPS relative to monoUb-PA was surprising since SARS-CoV-2 PLpro has been demonstrated to robustly process K48-linked polyUb chains and a preference for Ub over ISG15 (*27*, *28*).

**Fig. 4.**
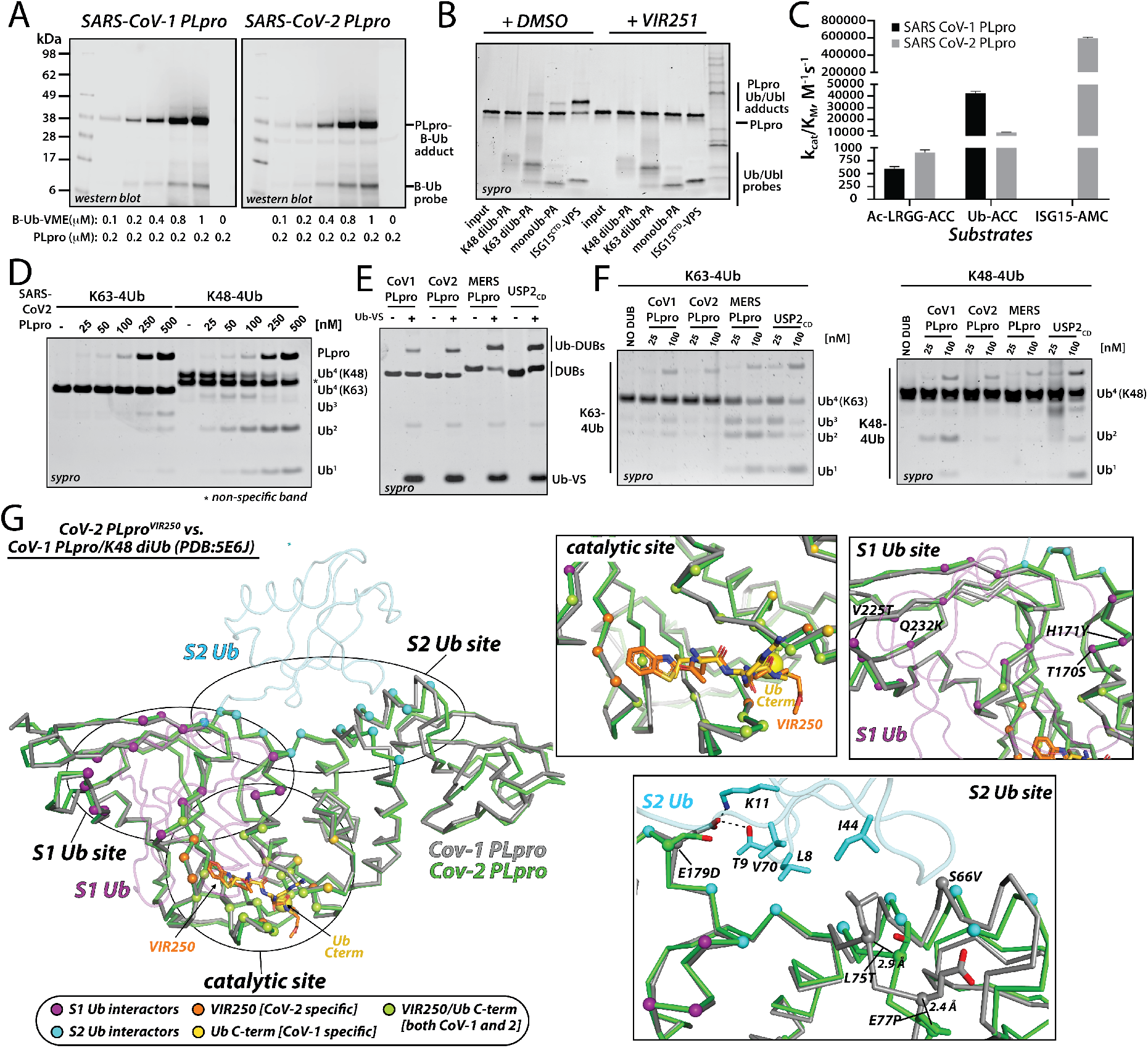
Processing of Ub and UbI variants by SARS-CoV-1 PLpro and SARS-CoV-2-PLpro. **(A)** SARS-CoV-1 PLpro and SARS-CoV-2-PLpro labelling by Biotin-Ub-VME. Recombinant enzymes were incubated with the indicated B-Ub-VME concentration for 45 min at 37 C, analyzed by SDS-PAGE, and subjected to western blot using fluorescent streptavidin Alexa Fluor 647 conjugate. **(B)** SARS-CoV-2-PLpro was pretreated with DMSO or VIR251 for 30 min at 37 C and subsequently incubated with the indicated activity based Ub/Ubl probe at RT for 2 min. This was followed by SDS-PAGE and sypro staining. **(C)** Kinetic parameters of selected substrates for SARS-CoV-PLpro and SARS-CoV-2-PLpro. Asterisk indicated that Cov-1 PLpro/ISG15-AMC pair were not analyzed in this study but Kcat/KM values have previously been determined. **(D)** The indicated tetreUb chains were incubated with CoV-2 PLpro for 30 mon at 37 C, subjected to SDS-PAGE and sypro staining. **(E)** Ubiquitin-Vinyl Sulfone labeling. Ub-VS was incubated for protease was incubated for 30 min at 37 C with the incubated protease, subjected to SDS-PAGE and sypro staining. **(F)** The incubated tetreUb chains were incubated with the indicated protease and analyzed as in D.

To more thoroughly examine these differences, we performed a comparison of the kinetics of SARS-CoV-1 and CoV-2 PLpro processing of LRGG-ACC, Ub-ACC, and ISG15-AMC fluorogenic substrates. The results of this experiment show that SARS-CoV-2 PLpro processes Ub-ACC 4-fold less efficiently compared to SARS-CoV-1 PLpro and that SARS-CoV-2 PLpro processes ISG15-AMC 60-fold more efficiently than Ub-ACC (Figure 4C). Further, SARS-CoV-2 PLpro, like SARS-CoV-1 PLpro, more robustly processes K48 tetraUb compared to K63 tetraUb (Figure 4D), and cross-links to the ABP Ub-VS similarly to SARS-CoV-1 PLpro, MERS PLpro, and the human DUB USP2_CD_ (Figure 4E). Yet, in side-by-side comparison, SARS-CoV-2 PLpro demonstrates a significantly diminished ability to process K48 tetraUb compared to SARS-CoV-1 PLpro (Figure 4F). This was an unexpected finding as we and others have shown before that SARS-CoV-1 PLpro displays a preference for recognition of K48 diUb linkages over ISG15 (*27*, *28*). As expected and shown before, both MERS PLpro and USP2_CD_ efficiently processes both types of Ub chains.

The significantly diminished ability of SARS-CoV-2 PLpro to process K48 polyUb chains compared to SARS-CoV-1 PLpro was surprising considering the very high overall similarity between the enzymes (83% identity, 9% similarity) (Figure 3D). To try to reconcile this apparent contradiction, we compared our SARS-CoV-2 PLpro structures with the previously reported structure of SARS-CoV-1-PLpro in complex with K48 diUb (Figure 4G) (*28*). This structure revealed three key interfaces: 1) the catalytic site that accommodates the C-terminus of Ub with L73, R74, G75, and G76 constituting the S4-S1 residues, 2) a binding site for the ‘S1 Ub’ which is the Ub N-terminal to the cleavage site in the K48 polyUb chain, and 3) a binding site for the ‘S2 Ub’ which is N-terminal to the S1 Ub (Figure 4G). Comparative analysis of the catalytic sites of SARS-CoV-1 and 2 PLpro shows a 100% conservation of residues involved in contacts to the S4-S1 positions of S1 Ub, VIR250, and VIR251 (Figure 3D) and unsurprisingly a very similar structure in this region (Figure 4G). While the Ub S1 site harbors more variability than the catalytic site the overall amino acid conservation is still very high (83% identity, 17% similarity) (Figure 3D) and the structures align well in this region (Figure 4G).

In contrast to the catalytic and S1 Ub sites, the S2 Ub site of SARS-CoV-2 PLpro harbors much less conservation at the amino acid level (67% identity, 13% similarity) compared to SARS-CoV-1 PLpro (Figure 3D), and there are several structural differences at these regions important for molecular recognition of the S2 Ub (Figure 4G). A key interaction surface at this interface is formed by the Ile44 hydrophobic patch (formed by Leu8, Ile44, and Val70), and in the SARS-CoV-1 PLpro/K48 diUb structure the Ile44 patch of the S2 Ub engages in a network of hydrophobic contacts with Leu75 (Figure 4G). Interestingly, this residue has changed to a threonine in SARS-CoV-2 which would be unable to engage in a similar network of contacts with S2 Ub as leucine. Further, much of the SARS-CoV-2 structure in proximity to the S2 Ub site adopts a slightly different structure including a ~3 Å translation of Thr75 of SARS-CoV-2 PLpro relative to Leu75 of SARS-CoV-1 PLpro and other notable amino acid changes of S66V and E77P (Figure 4G). Lastly, Glu179 of SARS-CoV-1 PLpro engages in hydrogen bonds to Thr9 and Lys11 of S2 Ub (Figure 4G). This residue has changed to an aspartate in SARS-CoV-2 PLpro, which is a conservative change, but the shorter aspartate side chain is unable to engage in a similar set of contacts (Figure 4G). Based on our analysis, we posit that the diminished ability of SARS-CoV-2 PLpro to process K48 polyUb is largely due to the aforementioned differences at the S2 Ub binding site. Consistent with this hypothesis, mutation of Leu75 of SARS-CoV-1 PLpro to serine resulted in a five-fold reduction in binding of K48 diUb with no apparent effect on monoUb (*27*). In combination with other changes in SARS-CoV-2 PLpro such as E179D, there appears to be a cumulative effect of several relatively minor changes between SARS-CoV-1 and 2 PLpro at the S2 binding site that altogether have a significant effect on their ability to process K48 polyUb. Whether these changes also account for the apparent preference of SARS-CoV-2 PLpro for ISG15 over Ub and whether these intriguing differences in the function of SARS-CoV-1 and 2 PLpro have any effect on the biology of the viruses remains to be seen.

## Discussion and Conclusions

The outbreak of the current coronavirus pandemic leading to COVID-19 disease has dramatically accelerated research into effective drugs and a vaccine to treat this disease. The genome of severe acute respiratory syndrome coronavirus 2 (SARS-CoV-2) consists of 29811 nucleotides that encode 29 proteins, two of which are proteases. The first of these, SARS-CoV-2-Mpro is used by the virus in the process of protein maturation. Its structure has already been described recently (*29*). The results of retargeting about 12,000 drugs and other leading structures resulted in selection of several candidates for further studies (*30*). To date, there is no information about the activity of the second protease, namely SARS-CoV-2-PLpro. We were hypothesizing that this enzyme, similarly to SARS-CoV-PLpro, in addition to participating in the process of virus protein maturation, also performs an additional function, which is to help the virus in evasion of the host innate immune responses by controlling the deubiquitination and deISGylation process. Thus, SARS-CoV-2-PLpro is also an excellent candidate for a drug, not only blocking virus replication, but also inhibiting the dysregulation of signaling cascades in infected cells (*9*). Knowledge of substrate preferences is equally important with understanding the structure of the protein, as it enables rational design of inhibitors or research on drug retargeting.

In our research, we decided to examine the SARS-CoV-2-PLpro substrate preferences at positions P4-P2 and compare them directly with the well-known SARS virus 2002/03 protein, SARS-CoV-PLpro. For this purpose, we used positional scanning technology using natural and unnatural amino acids (HyCoSuL). Library screening revealed that both enzymes recognize only Gly in P2 and possess broad in P3 and rather narrow substrate specificity at the P4 position. Moreover, direct analysis of the preferences of both enzymes demonstrate that the architecture of S4-S2 pockets is almost identical, because they recognize natural and unnatural amino acids practically in a very similar way. The differences in activity for a given amino acid between the two enzymes observed in some positions are very small, and there are no amino acids that are recognized by one enzyme only. This is also confirmed by the analysis of amino acids building S4-S2 pockets in both enzymes, which is identical (Figure 1B and S1). This is critically important information in the aspect of using information from research on inhibitors or retargeting of drugs conducted in the past for SARS-CoV-PLpro for immediate application to SARS-CoV-2-PLpro. Analysis of kinetic parameters for tetrapeptide substrates for both enzymes shows a high degree of similarity in terms of k_cat_/K_m_ values, proving that the catalytic yields of both enzymes are also similar. Importantly, the sequences containing unnatural amino acids at P4-P3 positions were recognized only by both SARS-PLpro, not MERS-PLpro and the human DUB UCH-L3.

We next leveraged the information we gained regarding the molecular rules governing substrate selectivity by SARS-CoV-2-PLpro to develop covalent inhibitors VIR250 and VIR251. These inhibitors proved to be active and selectively inhibited the SARS-CoV-1 and 2 PLpro, but exhibited much weaker activity towards MERS-PLpro and practically no activity towards human UCH-L3. This is excellent information in terms of conducting research towards the search for peptide antiviral compounds targeted to this enzyme. Importantly, our crystal structures of VIR250 and VIR251 in complex with SARS-CoV-2-PLpro reveals their inhibitory mechanisms and provides a structural basis for the observed substrate specificity profiles. Further, the unexpected finding that the P4 amino acids of VIR250 and VIR251 occupy opposite sides of the broad S4 pocket of SARS-CoV-2 PLpro and that there are additional regions of this pocket that are unengaged by either inhibitor raise the possibility that our structures will inform future drug discovery efforts aimed at generating more potent inhibitors.

Lastly, we decided to examine processing of Ub and Ubl variants by SARS-CoV-1 and 2 PLpro and found that SARS-CoV-2-PLpro harbors deISGylating activities similar to SARS-CoV-1-PLpro but its ability to hydrolyze K48-linked Ub chains is significantly diminished. This was an unexpected result considering the very high sequence identity between SARS-CoV-1 and 2 PLpro, however our structure analysis revealed subtle structural and sequence variations in the S2 Ub binding site of SARS-CoV-2 PLpro which we posit collectively diminish the ability of the S2 Ub of K48 polyUb to bind and subsequently be processed. Further, analysis of the enzyme kinetics of the Ub-ACC substrate indicates that it is efficiently processed by the enzyme, but the difference between the tetrapeptide substrate and ubiquitin is only about ten times, when in the case of SARS-CoV-1 PLpro this difference is around sixty times (Figure 4C). This indicates some differences between both enzymes in the aspect of interaction in the exosite binding region related to amino acids identity and similarity. Given the role of Ub and ISG15 conjugation in evasion of the host innate immune responses whether these intriguing differences in the function of SARS-CoV-1 and 2 PLpro have any effect on the biology of the viruses remains to be seen and will be the topic of future studies.

Collectively our work has revealed the molecular rules governing PLpro substrate specificity and reveals a very high level of sequence and structural similarity between SARS-CoV-1 and 2 PLpro in the substrate binding pocket. These findings signal that previously discovered information on SARS-CoV-1 PLpro can immediately be applied to the search for effective antiviral molecules as well as retargeting of known drugs for the inhibition of SARS-CoV-2 PLpro. Further, structures of the novel inhibitors VIR250 and VIR251 in complex with SARS-CoV-2 PLpro provides a framework for rational development of inhibitors with improved potency. Altogether, our data also gives a hope for design of a drug that can act as a pan-selective inhibitor against both SARS-CoV-PLpro and SARS-CoV-2-PLpro, and may have some universal value against emerging coronaviruses in the near future.

## Acknowledgments

This project was supported by the National Science Center grant 2015/17/N/ST5/03072 (Preludium 9) in Poland (W.R.) and the “TEAM/2017-4/32” project, which is carried out within the TEAM program of the Foundation for Polish Science, co-financed by the European Union under the European Regional Development Fund (M.D.). W.R. is a beneficiary of a START scholarship from the Foundation for Polish Science. This work is based upon research conducted at the Northeastern Collaborative Access Team beamlines, which are funded by the National Institute of General Medical Sciences from the National Institutes of Health (P30 GM124165). This research used resources of the Advanced Photon Source, a U.S. Department of Energy (DOE) Office of Science User Facility operated for the DOE Office of Science by Argonne National Laboratory under Contract No. DE-AC02-06CH11357. The X-ray crystallography facility used for this work is supported by the Office of the Vice President for Research at the Medical University of South Carolina. The liquid handling robot used was purchased via an NIH Shared Instrumentation Award (S10 RR027139-01). Research reported in this publication was supported by the NIH R01 GM115568 (S.K.O.), ES025166 (T.T.H.), and GM099040 (S.J.S.). Z.L. is a Hollings Cancer Center Postdoctoral Fellow, and S.P. is an American Cancer Society Postdoctoral Fellow (PF-18-235-01-RMC). The content of this study is solely the responsibility of the authors and does not necessarily represent the official views of the NIH.

## Data Availability

Atomic coordinates and structure factors are deposited in the RCSB with accession codes 6WUU (SARS-CoV-2 PLpro/VIR250) and 6WX4 (SARS-CoV-2 PLpro/VIR251).

## Competing interest

F.E.O. declares competing financial interests as co-founders and shareholder of UbiQ Bio BV. M.B. is an employee and shareholder of Arvinas, Inc. The remaining authors declare no competing interests.

## Author contributions

W.R. M.B., S.K.O., and M.D. conceived the project; M. D., W.R. T.T.H., S.K.O., and M.Z. designed the research; W.R., M.Z., Z.L., S.P. and D.N. performed the research and collected data; D.N., Z.L., S.J.S., M.B. and T.T.H. contributed enzymes; M.D., W.R., T.T.H., M.B., S.K.O., and M.Z. analyzed and interpreted the data; Z.L., and S.K.O. performed structural experiments including crystallization, X-ray data collection/processing, model building/refinement, and structural analyses; W.R., S.K.O, and M.D. wrote the manuscript and all authors critically revised the manuscript.

## Notes

### Summary of Updates

This version is updated with crystal structures of inhibitors with SARS-CoV-2-PLpro and biochemistry data.

