## Supplemental for "Activity profiling and structures of inhibitor-bound SARS-CoV-2-PLpro protease provides a framework for anti-COVID-19 drug design"

[6]Independent Consultant

[7]present address: Department of Biochemistry & Structural Biology University of Texas Health Science Center at San Antonio, San Antonio, TX, 78229 USA

[8]present address: Arvinas, Inc., 5 Science Park, New Haven, CT, 06511, USA

[9]equal contribution

**Content of Supporting Information**

1. Materials and Methods…………………………………………………..……….S3-S8

2. Supplemental Figure 1…………………………………...………..…..………….S9

3. Supplemental Figure 2…………………………………………………….……...…S10

4. Supplementary Table 1……………………………………………………..S11-S18

5. Supplementary Table 2…………………………………………………………..S19

6. Analysis data of synthesized chemical compounds…………….…..……...……..S20-S29

### Materials and Methods

#### Plasmids

The cDNA for Papain-Like protease (PLpro) corresponding to aa745-1061 of SARS-CoV-2 NSP3 was codon optimized for *E. coli* expression, synthesized and cloned into pGEX6P-1 (GE Healthcare, UK) using the BamHI and NotI sites by GeneUniversal (USA). The plasmid was transformed into BL21 (DE3) codon plus *E. coli* strain for protein expression.

#### Protein expression and purification

SARS-CoV-PLpro, UCH-L3 and MERS-PLpro were obtained as described earlier (*1, 2*). SARS-CoV-2-PLpro transformed cells cells were grown in LB broth at 37°C with shaking until the OD600 reached 1.5. 0.1 mM IPTG and 0.1 mM $ZnSO_4$ were added to induce protein expression overnight at 18°C. Cell pellet was resuspended in lysis buffer (20 mM Tris-Cl pH 8.0, 350 mM NaCl and 2 mM β-mercaptoethanol) and lysed using sonication. The lysate was cleared by centrifugation at 35000 g for 30 mins at 4°C. The lysate was passed onto Glutathione Sepharose 4B (GE) followed by washing with lysis buffer. The GST tagged PLpro was eluted in lysis buffer supplemented with 20 mM reduced Glutathione (pH 8.0). The fusion protein was cleaved using GST-PreScission protease at 4°C overnight followed with desalting and passing through fresh glutathione beads to remove cleaved GST and PreScission protease. The sample was further purified using Superdex 200 pg size-exclusion columns (GE) equilibrated with 20 mM Tris-Cl pH 8.0, 40 mM NaCl and 2 mM DTT. The purified protein was then concentrated to ~10 mg/ml and snap frozen in liquid nitrogen for later use.

#### Reagents

The reagents used for the solid-phase peptide synthesis (SPPS) were as follows: Rink Amide (RA) resin (particle size 100-200 mesh, loading 0.74 mmol/g), 2-chlorotrityl chloride resin (particle size 100-200 mesh, loading 0.97 mmol/g), all Fmoc-amino acids, *O*-benzotriazole-*N*,*N*,*N*`,*N*`-tetramethyl-uronium-hexafluoro-phosphate (HBTU), 2-(1-H-7-azabenzotriazol-1-yl)-1,1,3,3-tetramethyluranium hexafluorophosphate (HATU), piperidine, diisopropylcarbodiimide (DICI) and trifluoroacetic acid (TFA), purchased from Iris Bioteh GmbH (Marktredwitz, Germany); anhydrous *N*-hydroxybenzotriazole (HOBt) from Creosauls Louisville, KY, USA; 2,4,6-collidine (2,4,6-trimethylpyridine), HPLC-grade acetonitrile, triisopropylsilane (TIPS), *t*Bu-N-allyl carbamate, toluene, methyl acrylate, dichlorophenylborane and 2nd generation Grubbs catalyst from Sigma-Aldrich (Poznan, Poland); and *N,N*-diisopropylethylamie (DIPEA) from VWR International (Gdansk, Poland). *N,N*-dimethylformamide (DMF), dichloromethane (DCM), methanol (MeOH), diethyl ether ($Et_2O$), acetic acid (AcOH), and phosphorus pentoxide ($P_2O_5$), obtained from Avantor (Gliwice, Poland). Individual substrates, Ub-ACC and B-Ub-VME were purified by HPLC on a Waters M600 solvent delivery module with a Waters M2489 detector system using a semipreparative Wide Pore C8 Discovery column and Jupiter 10 µm C4 300 Å column (250 x 10 mm). The solvent composition was as follows: phase A (water/0.1% TFA) and phase B (acetonitrile/0.1% TFA). The purity of each compound was confirmed with an analytical HPLC system using a Jupiter 10 µm C4 300 Å column (250 x 4.6 mm). The solvent composition was as follows: phase A (water/0.1% TFA) and phase B (acetonitrile/0.1% TFA); gradient, from 5% B to 95% B over a period of 15 or 20 min. The molecular weight of each substrate and B-Ub-VME was confirmed by high-resolution mass spectrometry on a High-Resolution Mass Spectrometer Waters LCT premier XE with electrospray ionization (ESI) and a time-of-flight (TOF) module.

**Combinatorial and defined substrate library synthesis.**

Detailed protocol of combinatorial and defined tetrapeptide fluorogenic substrate library synthesis was described elsewhere (*3*).

**Determination of SARS-CoV and SARS-CoV-2-PLpro substrate specificity**

Library screening was performed using a spectrofluorometer (Molecular Devices Spectramax Gemini XPS) in 96-well plates containing substrates and enzymes. Assay conditions were 1 µL of substrate in DMSO and 99 µL of enzyme, which had been incubated for 15 min at 37°C in assay buffer (150 mM NaCl, 20 mM Tris, 5 mM DTT, pH 8.0 for SARS-CoV PLpro; 5 mM NaCl, 20 mM Tris, 5 mM DTT, pH 8.0 for SARS-CoV-2 PLpro). The final substrate concentration in each well was 200 µM of combinatorial library and 100 µM of defined P2 library. The final enzyme concentration was 1 µM SARS-CoV-PLpro and 0.5 µM SARS-CoV-2-PLpro for P3 and P4 sublibraries and 0.1 µM SARS-CoV-PLpro and 75 nM SARS-CoV-2-PLpro for Ac-Leu-Arg-P2-Gly-ACC. The release of ACC was measured continuously for 45 min ($\lambda_{ex}$ = 355 nm, $\lambda_{em}$ = 460 nm). SARS-CoV and SARS-CoV-2-PLpro substrate specificity profiles were established by setting the highest relative fluorescence unit per second (RFU/s) for the best substrate as to 100% and adjusting other results accordingly.

**Synthesis of tetrapeptide fluorogenic substrates and Ub-ACC**

Individual fluorogenic substrates were synthesized on a solid support using the SPPS method as previously described (*4, 5*). Each substrate was purified by HPLC and analyzed using analytical HPLC and HRMS. The purity of each compound was ≥95%. The individual substrates were dissolved at 20 mM in DMSO and stored at -80°C until use.

**Kinetic studies of individual tetrapeptide substrates and Ub-ACC**

Individual substrate hydrolysis was measured in the same assay conditions as for library screening. The final substrate concentration was 10 μM, SARS-CoV and SARS-CoV-2 PLpro concentration was 0.1 μM, MERS-CoV PLpro was 2.5 μM, and UCH-L3 was 8.8 μM. MERS-CoV PLpro and UCH-L3 were incubated for 30 min at 37°C in assay buffer (MERS-CoV PLpro: 150 mM NaCl, 20 mM Tris, 5 mM DTT, pH 8.0; UCH-L3: 50 mM HEPES, 0.5 mM EDTA, 5 mM DTT, pH 7.5) prior to add into the wells on plate. The measurements were repeated at least three times and the results were presented as mean values with standard deviations. Kinetic parameters were determined for selected tetrapeptide substrates and Ub-ACC toward SARS-CoV and SARS-CoV-2 PLpro. Wells contained 20 μL of substrate in assay buffer at eight different concentrations (0.88 μM to 20 μM) and 80 μL of enzyme (0.5 μM SARS-CoV and SARS-CoV-2 PLpro for tetrapeptide substrates and 80 nM SARS-CoV-2 PLpro and 10 nM SARS-CoV PLpro for Ub-ACC). Substrate hydrolysis was measured for 30 min at the appropriate wavelength ($\lambda_{ex}$ = 355 nm, $\lambda_{em}$ = 460 nm). Each experiment was carried out at least three times and the results reported as averages with standard deviation. Due to the precipitation of tetrapeptide substrates at high concentrations, only the specificity constant ($k_{cat}/K_M$) was determined. When $[S_0] << K_M$, the plot of $v_i$ (the initial velocities) versus $[S_0]$ yields a straight line with slope representing $V_{max}/K_M$, $k_{cat}/K_M$ = slope/E (E – total enzyme concentration). Kinetic parameter for ISG15-AMC toward SARS-CoV-2 PLpro was determined in the same manner as described above. The final enzyme concentration was 1 nM and the final substrate concentration was ranging from 0.3 μM to 5 μM.

**SARS-CoV and SARS-CoV-2-PLpro inhibitor and B-Ub-VME synthesis**

Inhibitor synthesis was performed in the three sequential stages. In the first step vinyl methyl ester as a reactive group was synthesized according to published protocol (*6*). *t*Bu-N-allyl carbamate (500 mg, 3.2 µmol) was dissolved in 10 mL of anhydrous toluene. Methyl acrylate (580 µL, 6.4 µmol), dichlorophenylborane (42 µL, 0.32 µmol) and 2nd generation Grubbs catalyst (50 mg) were added. The reaction was carried out under reflux at 40°C with stirring overnight. After 12 h, the solvent was evaporated under reduce preasure, and the mixture was purified by column chromatography on silica gel (Hex/EtOAc 5:1). The crude product was obtained as a yellowish oil. *t*Bu group deprotection was performed by adding TFA/DCM/TIPS (4.2 mL, 3/1/0.2, *v/v/v*) cleavage mixture for 45 min with stirring. TFA*$H_2N$-Gly-VME was then crystallized in cold $Et_2O$ and stored at -20°C. In the second step Ac-P4-P3-Gly-OH fragments were synthesized using 2-chlorotrityl chloride resin as previously described (*7*). In the last step Ac-P4-P3-Gly-OH fragment (1.2 eq) was coupled to the reactive group (1 eq) using HATU (1.2 eq) and 2,4,6-collidine (3 eq) as a coupling reagents in DMF. The reaction was carried out at RT with stirring for 2h. The reaction mixture was diluted in ethyl acetate, washed once with 5% citric acid, once with 5 % $NaHCO_3$ and once brine, dried over $MgSO_4$, and concentrated under reduce pressure. To remove side chain amino acid protecting groups, Ac-P4-P3-Gly-Gly-VME was added to a mixture of TFA/DCM/TIPS (% v/v/v, 70:27:3). After 30 min, solvents were removed and inhibitor was purified on HPLC. B-Ub-VME was synthesized according to synthetic protocol described elsewhere (*8, 9*).

**Determination of DUB inhibition**

To assess activity and selectivity of designed SARS-CoV and SARS-CoV-2-PLpro inhibitors DUBs were incubated with inhibitors at eight different concentrations (2.3 µM to 300 µM) for 30 min at 37°C in assay buffers. DUB residual activity was estimated using Ac-Leu-Arg-Gly-Gly-ACC (50 µM). Assay conditions were 20 µL of inhibitor, 60 µL of DUB (0.3 µM SARS-CoV PLpro, 0.1 µM SARS-CoV-2 PLpro, 2.5 µM MERS-CoV PLpro, and 8 µM UCH-L3), and 20 µL of substrate (50 µM). Inhibition assays were measured for 40 min and repeated at least three times. The results were established as mean values with standard deviations.

**Crystallization**

3 uM SARS-CoV-2-PLpro was reacted with 30 uM peptide inhibitor in 5 mM NaCl, 20 mM Tris HCl pH 8.0 at 37°C for 20 min. Protein was concentrated using a 30 kD cutoff Amicon Ultra Filter and desalted into 5 mM NaCl, 20 mM Tris HCl pH8.0, 10 mM DTT. Final protein concentration was 5-10 mg/ml. VIR250 complex crystals were grown by mixing 0.4 ul protein and 0.4 ul well solution containing 0.2 M Lithium Ammonium tribasic, 20% PEG3350 on a 96 well sitting plate at 18°C. Crystals were cryo protected by well solution plus 25% (v/v) ethylene

glycol and snap frozen in liquid nitrogen. VIR251 complex crystals were grown by mixing 0.2 ul protein sample with 0.2 ul well solution containing 0.8 M Potassium sodium tartrate tetrahydrate, 0.1 M Tris HCl pH 8.5 and 0.5% w/v Polyethylene glycol monomethyl ether 5,000 on a 96 well sitting plate at 18 degrees. Crystals were cryo protected by 25% (v/v) ethylene glycol and flash frozen with liquid nitrogen.

**Structure determination and refinement**

A complete data set was collected from the SARS-CoV-2-PLpro/VIR250 crystals to 2.79 Å resolution at the Advanced Photon Source (APS), NE-CAT beamline 24-IDC at a wavelength of 0.979 Å. Data set was indexed, integrated, and scaled using HKL2000. Crystal belongs to space group $P2_1$ with unit cell dimensions $a$ = 58.4, $b$ = 189.7, $c$ = 63.1, β=98.7°. There are four SARS-CoV-2-PLpro/VIR250 complexes per asymmetric unit. The structure was solved by molecular replacement using the program PHASER. The search model was apo SARS-CoV-2-PLpro structure (PDB: 6W9C). Apparent ligand density for both Fo-Fc and 2Fo-Fc maps was observed projecting off Cys111 after first round of refinement. Model and restraints for VIR250 was prepared using Phenix.Elbow. Model of SARS-CoV-2-PLpro/VIR250 was subjected to iterative rounds of refinement and rebuilding using PHENIX (*10*) and COOT (*11*).

For SARS-CoV-2-PLpro/VIR251 crystals, data was collected and processed as described above for VIR250, to a resolution of 1.65 Å. The crystal belongs to space group I222 with unit cell dimensions $a$ = 44.9, $b$ = 113.5, $c$ = 151.1. There is one SARS-CoV-2-PLpro/VIR251 complex per asymmetric unit. The structure was determined by molecular replacement with Phaser and the search model was SARS-CoV-2-PLpro/VIR250 structure above (PDB: 6WUU). Structure with ligand was refined as described above for the VIR250 structure.

The final two models for Plpro-VIR250 and Plpro-VIR251 complexes have R/Rfree values of 0.195/0.230 and 0.170/0.196, respectively. The two structures also have excellent geometry as assessed using Molprobity: favored (95.3%), allowed (4.6%), and outliers (0.1%) for the PLpro/VIR250 structure, and favored (97.0%), allowed (3.0%), and outliers (0%) for the PLpro/VIR251 structure.

**PLpro-Ub/Ubl ABP panel assay**

K48 diUb-PA, K63 diUb-PA, monoUb-PA and $ISG15^{CTD}$-VPS probes used in this experiment were generous gifts of UbiQ Bio. 3 uM Plpro was incubated with 30 uM inhibitor or DMSO at 37 for 20 min and put on ice. Reaction buffer contains 5 mM NaCl, 20 mM Tris HCl pH8.0.

Then the indicated Ub/Ubl ABPs were mixed with PLpro at 4.5 uM and 2.7 uM respectively at RT for 2 min. Reactions were terminated by adding SDS sample buffer, subjected to SDS-PAGE sypro staining.

**SARS-CoV and SARS-CoV-2-PLpro labelling by B-Ub-VME**

Enzymes (200 nM) were incubated with different B-Ub-VME concentrations (100, 200, 400, 800, and 1000 nM) in assay buffer (150 mM NaCl, 20 mM Tris, 5 mM DTT, pH 8.0 for SARS-CoV PLpro; 5 mM NaCl, 20 mM Tris, 5 mM DTT, pH 8.0 for SARS-CoV-2 PLpro) for 45 min at 37°C. Then 3x SDS/DTT was added, and the samples were boiled for 5 min at 95°C and resolved on 4-12% Bis-Tris Plus 12-well gels. Electrophoresis was performed at 200 V for 29 min. Next, the proteins were transferred to a nitrocellulose membrane (0.2 µm, Bio-Rad) for 60 min at 10 V. The membrane was blocked with 2% BSA in Tris-buffered saline with 0.1% (v/v) Tween 20 (TBS-T) for 60 min at RT. B-Ub-VME was detected with a fluorescent streptavidin Alexa Fluor 647 conjugate (1:10 000) in TBS-T with 1% BSA using an Azure Biosystems Sapphire Biomolecular Imager and Azure Spot Analysis Software.

**Gel-based Ub Chain Cleavage Assays and Ub-VS labeling**

Tetra-ubiquitin chains (K48- and K63-linked; BostonBiochem) were cleaved in a reaction volume of 10uL (in 20mM Tris pH=7.5, 150mM NaCl, 5mM DTT) with 25-500nM PLpro or USP2 catalytic domain (BostonBiochem), as indicated in figures. Ubiquitin-Vinyl Sulfone-labeling was performed in 10uL (in 20mM Tris pH=7.5, 150mM NaCl, 5mM DTT) with 1.5µM Ub-VS (BostonBiochem) and 0.25µM PLpro or USP2 catalytic domain. Reactions were incubated at 37˚C for 30min, terminated with sample loading buffer (4X LDS, Invitrogen), and analyzed by SDS-PAGE (4-12% Bis-Tris, NuPAGE) and SYPRO Ruby staining. Gels were imaged using an Azure biosystems c500 imager.

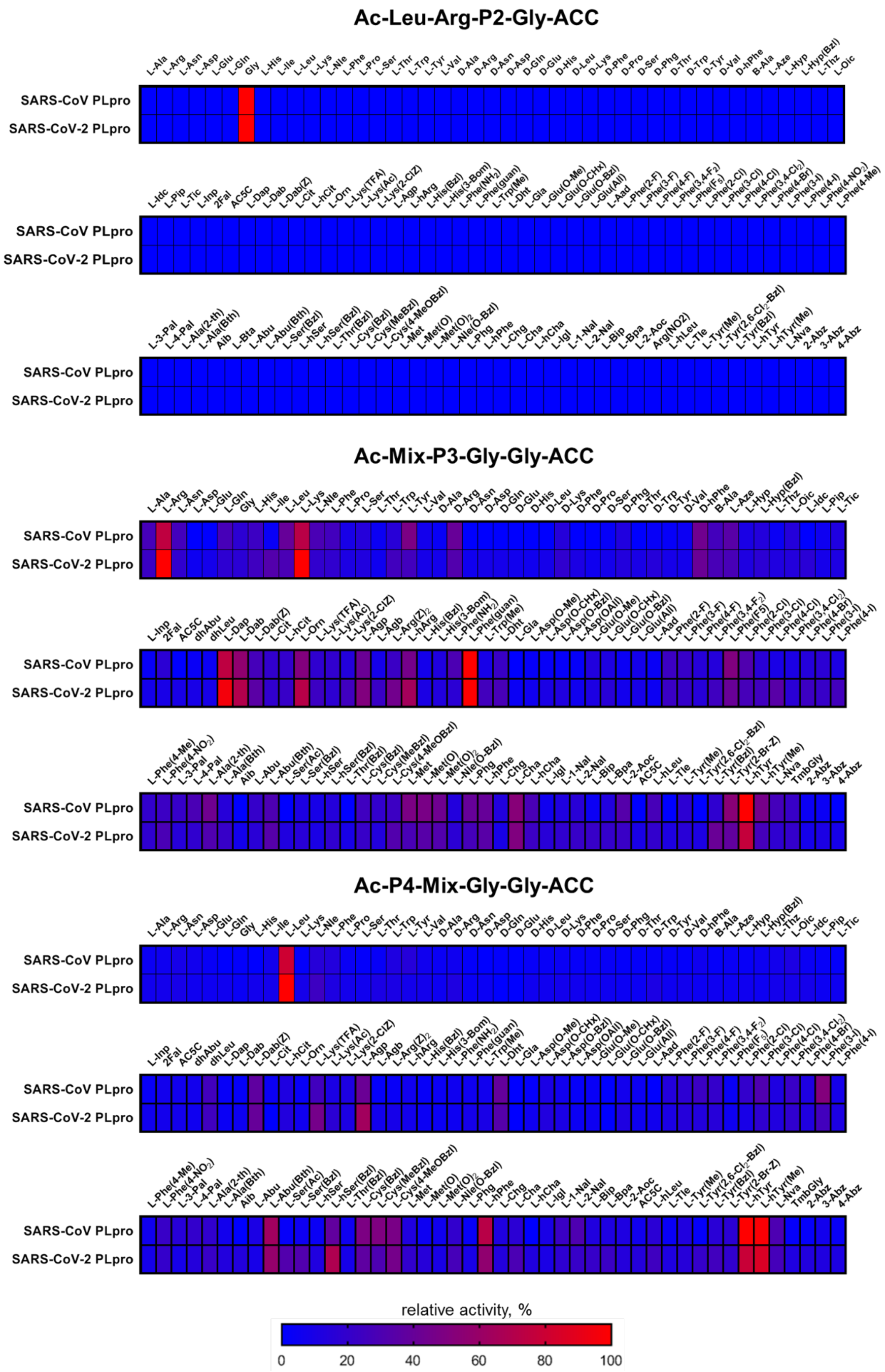

**Supplementary Figure 1. SARS-CoV-PLpro and SARS-CoV-2-PLpro substrate specificity profiles presented as heat maps.**

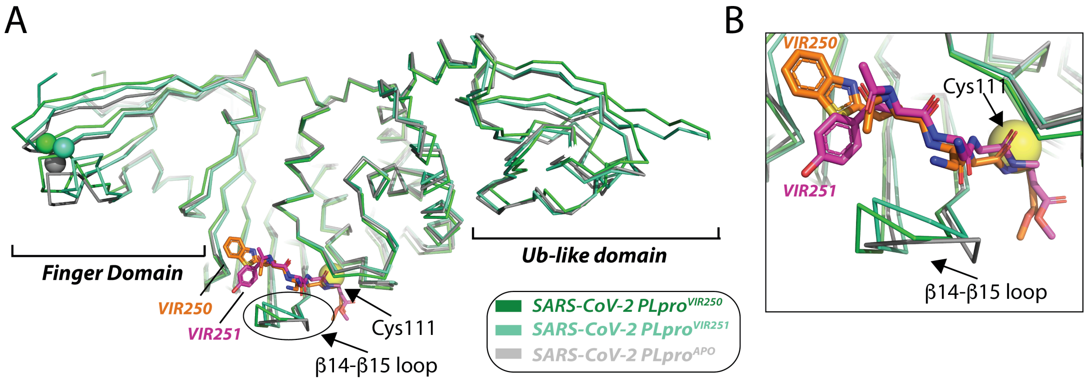

**Supplementary Figure 2. Comparision of apo versus VIR250 and VIR251 bound SARS-CoV-2 PLpro.** (**A**) The indicated structures were superimposed and are shown as ribbons. Zinc atoms are shown as spheres, colored according to their representative structure. Regions of conformational variability between apo and VIR250 and VIR251 structures are labeled including the Finger Domain, the Ub-like domain, and the β14-β15 loop. Catalytic cysteine 111 of SARS-CoV-2 PLpro is shown as sticks with its sg atom shown as yellow sphere. (**B**) Magnified view of the conformational differences around the β14-β15 loop of SARS-CoV-2 PLpro.

**Table S1**. Structures of fixed natural and unnatural amino acids used in combinatiorial library.

| No | Structure + code | No | Structure + code | No | Structure + code |
|---|---|---|---|---|---|
| 1 | 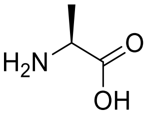 L-Ala | 2 | 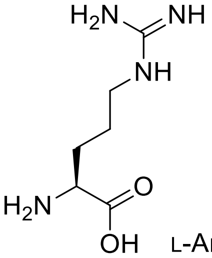 L-Arg | 3 | 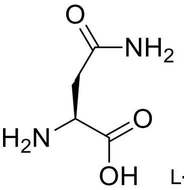 L-Asn |
| 4 | 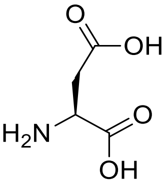 L-Asp | 5 | 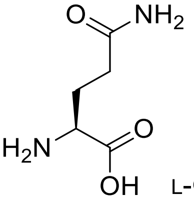 L-Gln | 6 | 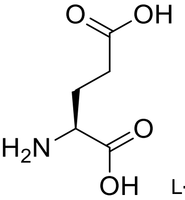 L-Glu |
| 7 | 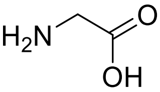 Gly | 8 | 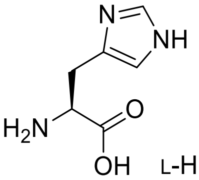 L-His | 9 | 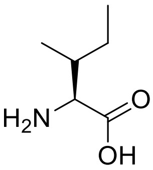 L-Ile |
| 10 | 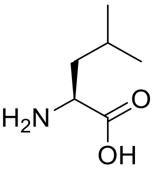 L-Leu | 11 | 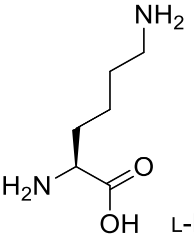 L-Lys | 12 | 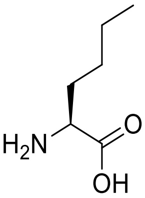 L-Nle |
| 13 | 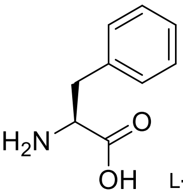 L-Phe | 14 | 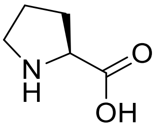 L-Pro | 15 | 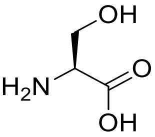 L-Ser |
| 16 | 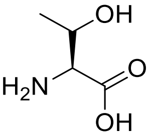 L-Thr | 17 | 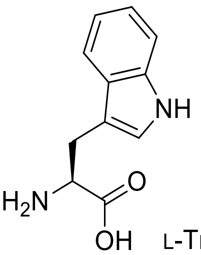 L-Trp | 18 | 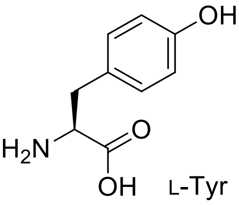 L-Tyr |

19 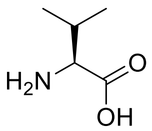 L-Val

20 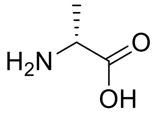 D-Ala

21 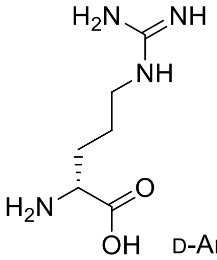 D-Arg

22 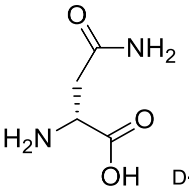 D-Asn

23 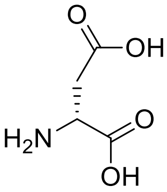 D-Asp

24 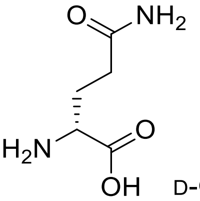 D-Gln

25 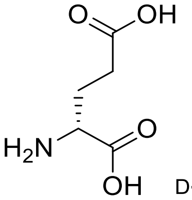 D-Glu

26 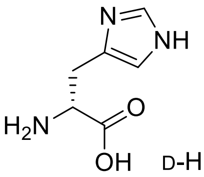 D-His

27 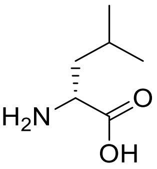 D-Leu

28 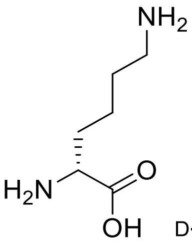 D-Lys

29 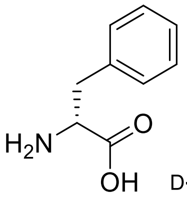 D-Phe

30 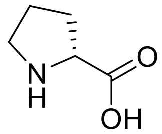 D-Pro

31 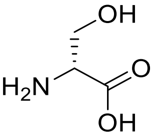 D-Ser

32 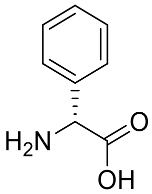 D-Phg

33 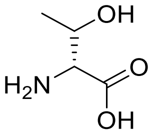 D-Thr

34 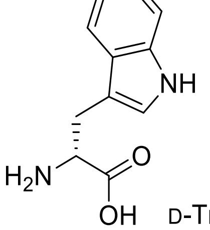 D-Trp

35 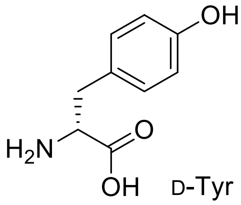 D-Tyr

36 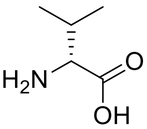 D-Val

37 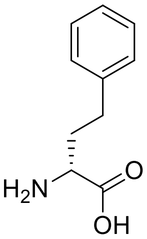 D-hPhe

38 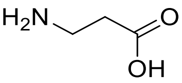 β–Ala

39 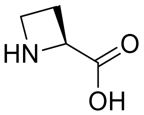 L-Aze

40 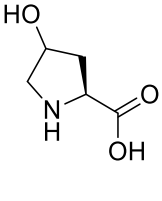 L-Hyp

41 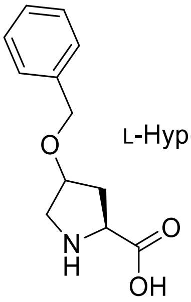 L-Hyp(Bzl)

42 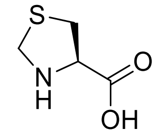 L-Thz

43 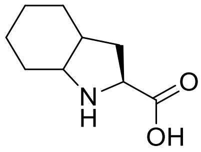 L-Oic

44 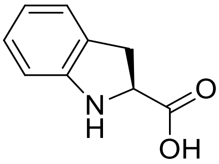 L-Idc

45 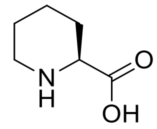 L-Pip

46 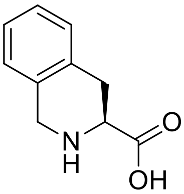 L-Tic

47 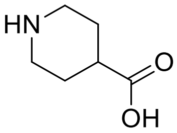 Inp

48 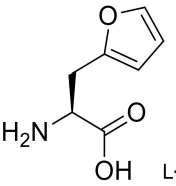 L-2Fal

49 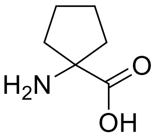 AC5C

50 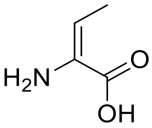 dhAbu

51 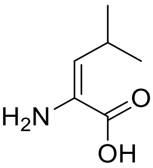 dhLeu

52 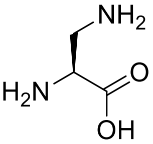 L-Dap

53 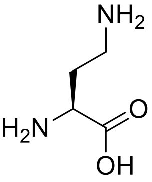 L-Dab

54 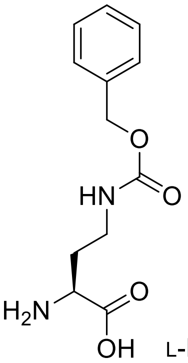 L-Dab(Z)

55 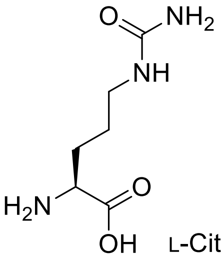 L-Cit

56 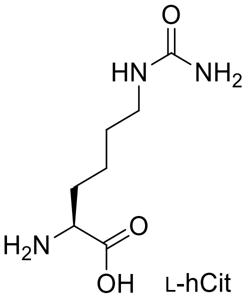 L-hCit

57 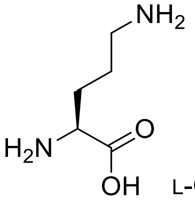 L-Orn

58 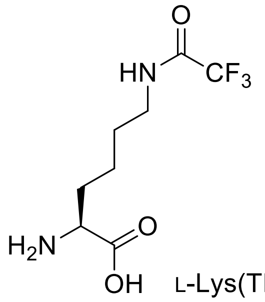 L-Lys(TFA)

59 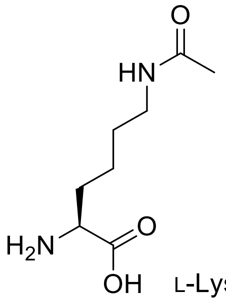 L-Lys(Ac)

60 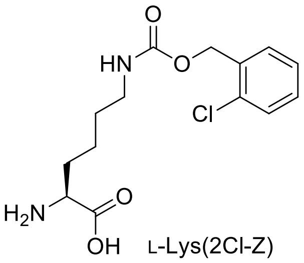 L-Lys(2Cl-Z)

61 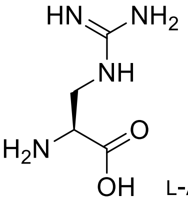 L-Agp

62 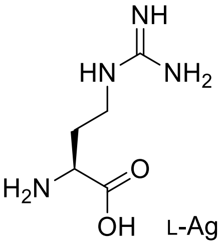 L-Agb

63 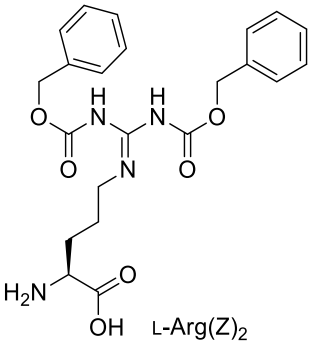 L-Arg(Z)$_2$

64 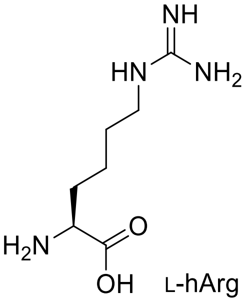 L-hArg

65 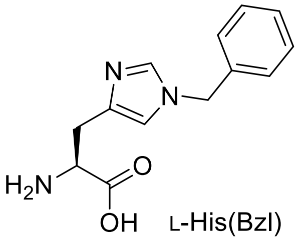 L-His(Bzl)

66 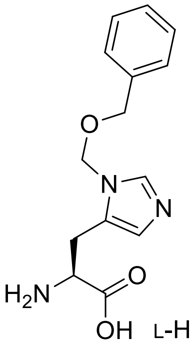 L-His(3-Bom)

67 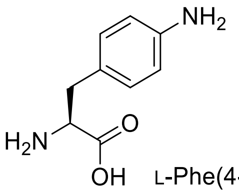 L-Phe(4-NH$_2$)

68 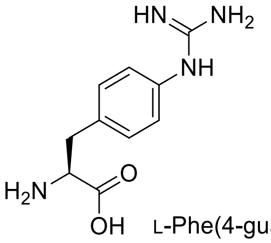 L-Phe(4-guan)

69 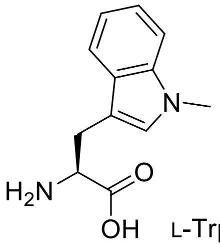 L-Trp(Me)

70 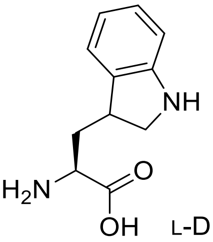 L-Dht

71 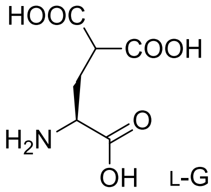 L-Gla

72 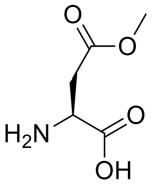 L-Asp(O-Me)

73 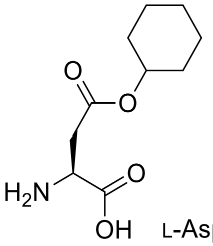 L-Asp(O-cHx)

74 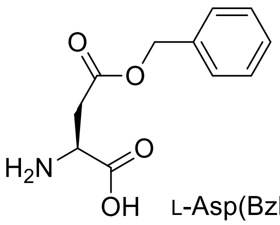 L-Asp(Bzl)

75 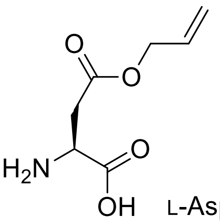 L-Asp(All)

76 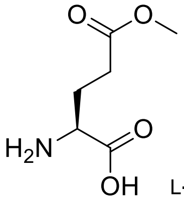 L-Glu(O-Me)

77 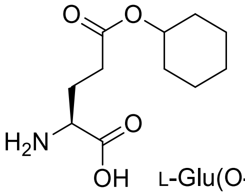 L-Glu(O-cHx)

78 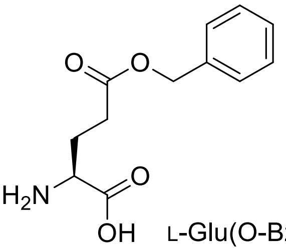 L-Glu(O-Bzl)

79 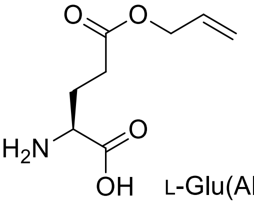 L-Glu(All)

80 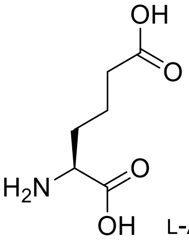 L-Aad

81 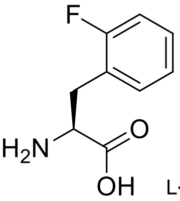 L-Phe(2-F)

82 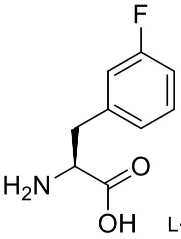 L-Phe(3-F)

83 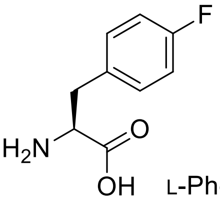 L-Phe(4-F)

84 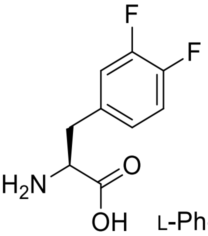 L-Phe(3,4-$F_2$)

85 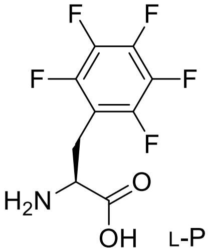 L-Phe($F_5$)

86 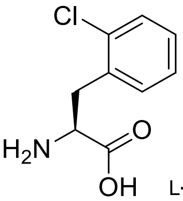 L-Phe(2-Cl)

87 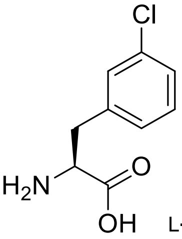 L-Phe(3-Cl)

88 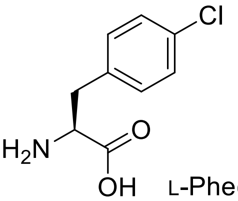 L-Phe(4-Cl)

89 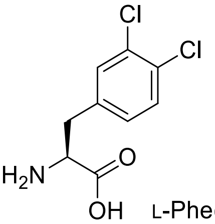 L-Phe(3,4-$Cl_2$)

90 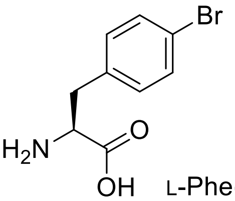 L-Phe(4-Br)

91 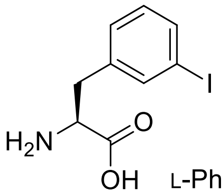 L-Phe(3-I)

92 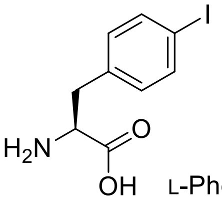 L-Phe(4-I)

93 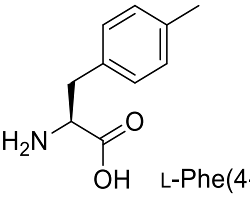 L-Phe(4-Me)

94 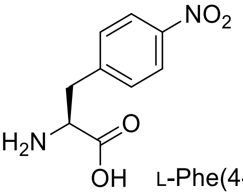 L-Phe(4-$NO_2$)

95 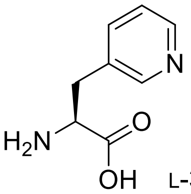 L-3-Pal

96 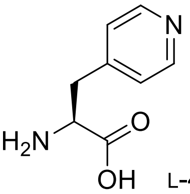 L-4-Pal

97 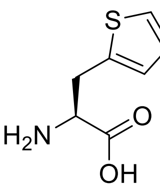 L-Ala(2-th)

98 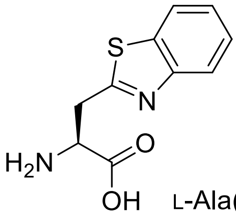 L-Ala(Bth)

99 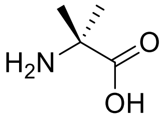 Aib

100 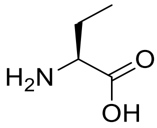 L-Abu

101 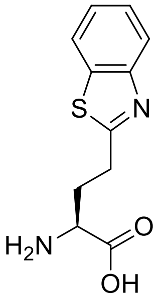 L-Abu(Bth)

102 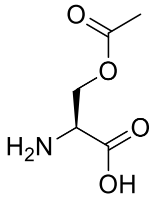 L-Ser(Ac)

103 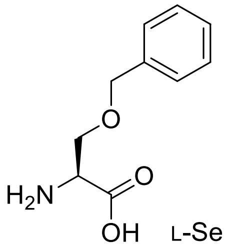 L-Ser(Bzl)

104 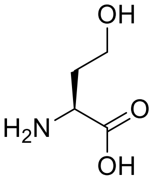 L-hSer

105 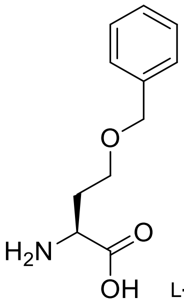 L-hSer(Bzl)

106 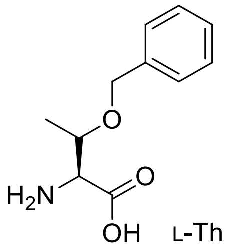 L-Thr(Bzl)

107 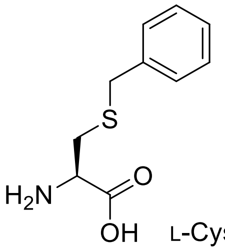 L-Cys(Bzl)

108 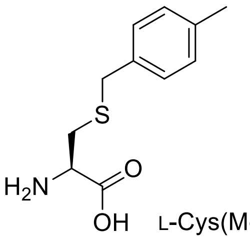 L-Cys(MeBzl)

109 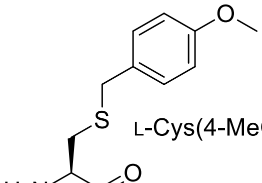 L-Cys(4-MeOBzl)

110 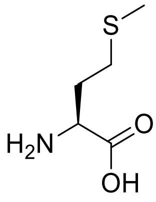 L-Met

111 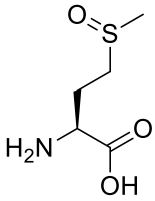 L-Met(O)

112 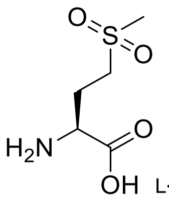 L-Met($O_2$)

113 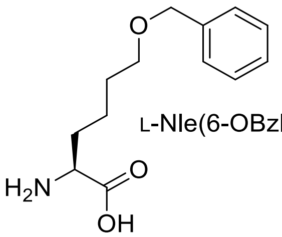 L-Nle(6-OBzl)

114 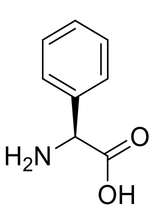 L-Phg

115 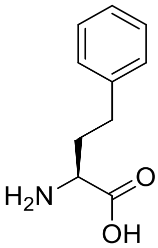 L-hPhe

116 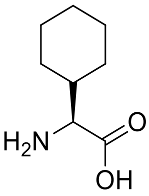 L-Chg

117 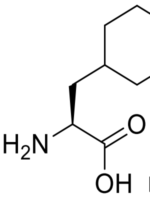 L-Cha

118 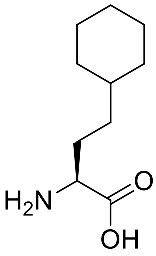 L-hCha

119 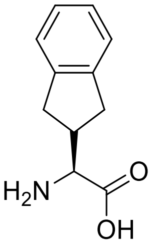 L-Igl

120 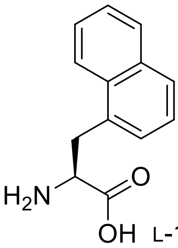 L-1-Nal

121 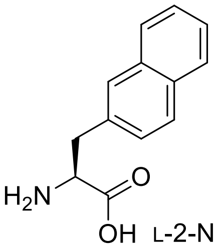 L-2-Nal

122 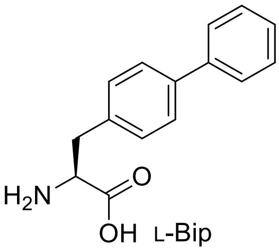 L-Bip

123 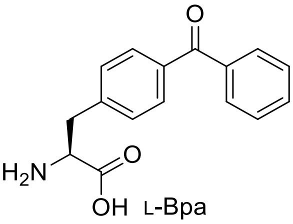 L-Bpa

124 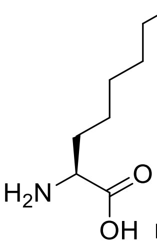 L-2-Aoc

125 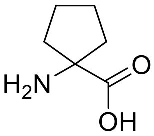 AC5C

126 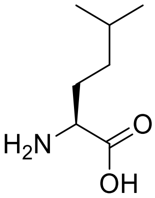 L-hLeu

127 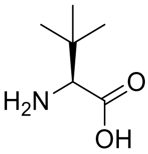 L-Tle

128 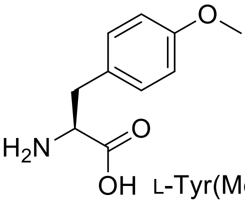 L-Tyr(Me)

129 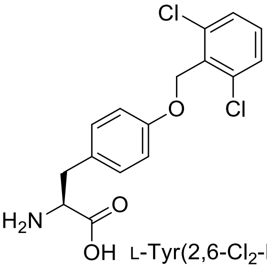 L-Tyr(2,6-$Cl_2$-Bzl)

130 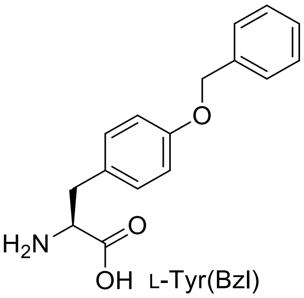 L-Tyr(Bzl)

131 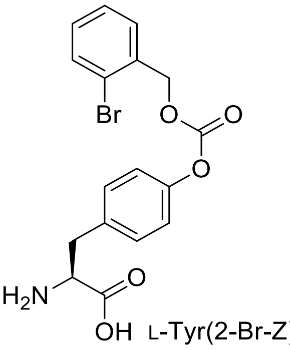 L-Tyr(2-Br-Z)

132 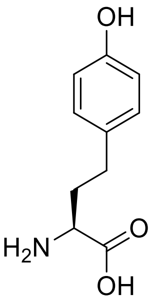 L-hTyr

133 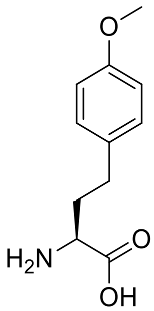 L-hTyr(Me)

134 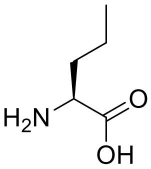 L-Nva

135 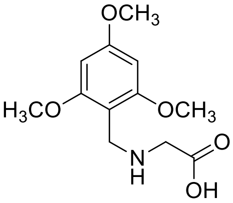 TmbGly

136 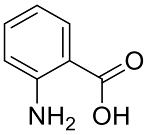 2-Abz

137 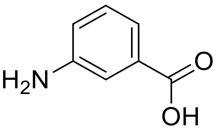 3-Abz

138 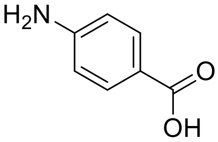 4-Abz

| | SARS-CoV-2 PLpro/VIR250 complex | SARS-CoV-2 PLpro/VIR251 complex |
|---|---|---|
| PDB ID | 6WUU | 6WX4 |
| Source | APS 24 IDC | APS 24 IDC |
| Wavelength (Å) | 0.979 | 0.979 |
| Resolution Limits (Å) | 94.9-2.79 (2.93-2.79) | 90.7-1.65 (1.68-1.65) |
| Space Group | $P2_1$ | I222 |
| Unit Cell (Å) *a*, *b*, *c* | 58.4, 189.7, 63.1 | 44.9, 113.5, 151.1 |
| Unit Cell (°) α, β, γ | 90, 98.7, 90 | 90, 90, 90 |
| Number of observations | 237255 | 611490 |
| Number of reflections | 33589 (4431) | 46470 (2152) |
| Completeness (%) | 99.6 (99.1) | 99.5 (91.0) |
| Mean I/σI | 10.1 (1.1) | 15.5 (1.2) |
| $CC_{1/2}$ | 0.996 (0.437) | 0.998 (0.531) |
| $R_{merge}$ [a] | 0.158 (1.79) | 0.094 (1.73) |
| $R_{pim}$ | 0.064 (0.753) | 0.027 (0.513) |
| Refinement Statistics | | |
| Resolution Limits (Å) | 94.9-2.79 (2.86-2.79) | 90.7-1.65 (1.70-1.65) |
| # of reflections (work/free) | 31489 (1992) | 44301 (2000) |
| Completeness (%) | 99.3 (98.0) | 99.5 (94.0) |
| Protein/solvent/ligand atoms | 10122/8/144 | 2545/194/35 |
| $R_{cryst}$ [b] | 0.195 (0.299) | 0.170 (0.241) |
| $R_{free}$ | 0.230 (0.333) | 0.196 (0.256) |
| Bonds (Å)/ Angles (°) | 0.004/0.737 | 0.006/0.844 |
| B-factors: protein/solvent/ligand ($Å^2$) | 81.2/84.8/78.2 | 37.7/43.5/30.6 |
| Ramachandran plot statistics (%) | | |
| favored | 95.3 | 97.0 |
| allowed | 4.6 | 3.0 |
| outliers | 0.1 | 0 |
| MolProbity score | 1.48- 100$^{th}$ percentile (N=4482, 2.79 Å ± 0.25Å) | 1.19- 99$^{th}$ percentile (N=789, 1.65 Å ± 0.25Å) |

Parentheses indicate statistics for the high-resolution data bin for x-ray data.
a. $R_{merge} = \sum hkl \sum i |I(hkl)i - \langle I(hkl)\rangle| / \sum hkl \sum i \langle I(hkl)i\rangle$.
b. $R_{cryst} = \sum hkl |Fo(hkl)-Fc(hkl)| / \sum hkl |Fo(hkl)|$, where *Fo* and *Fc* are observed and calculated structure factors, respectively.

**Table S2. Crystallographic Data and Refinement Statistics**

### LC-MS analysis of synthesized chemical compounds

### Ac-Leu-Arg-Gly-Gly-ACC

HRMS for $C_{29}H_{41}N_9O_8$ ($m/z_{calcd}$ = 644.3151; $m/z_{found}$ = 644.3158)

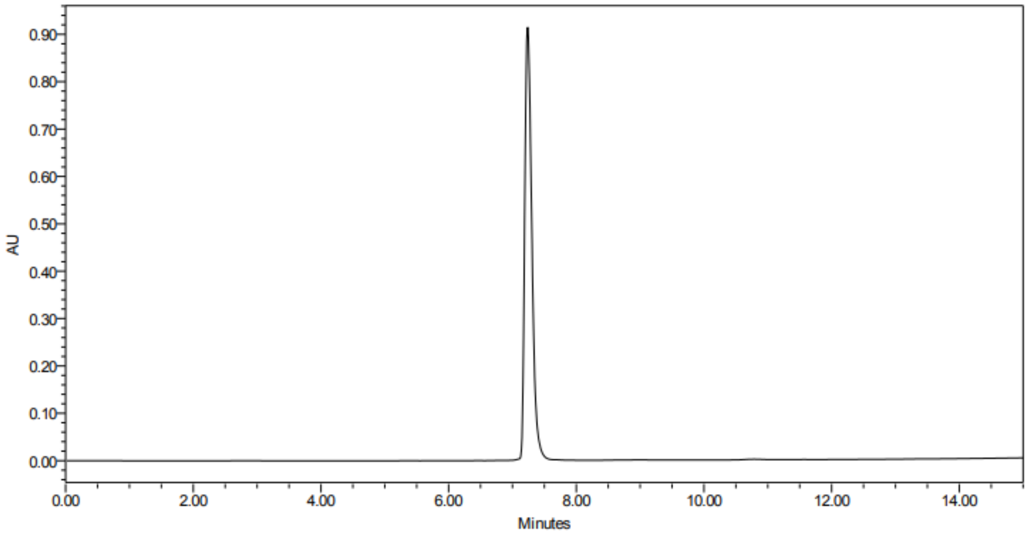

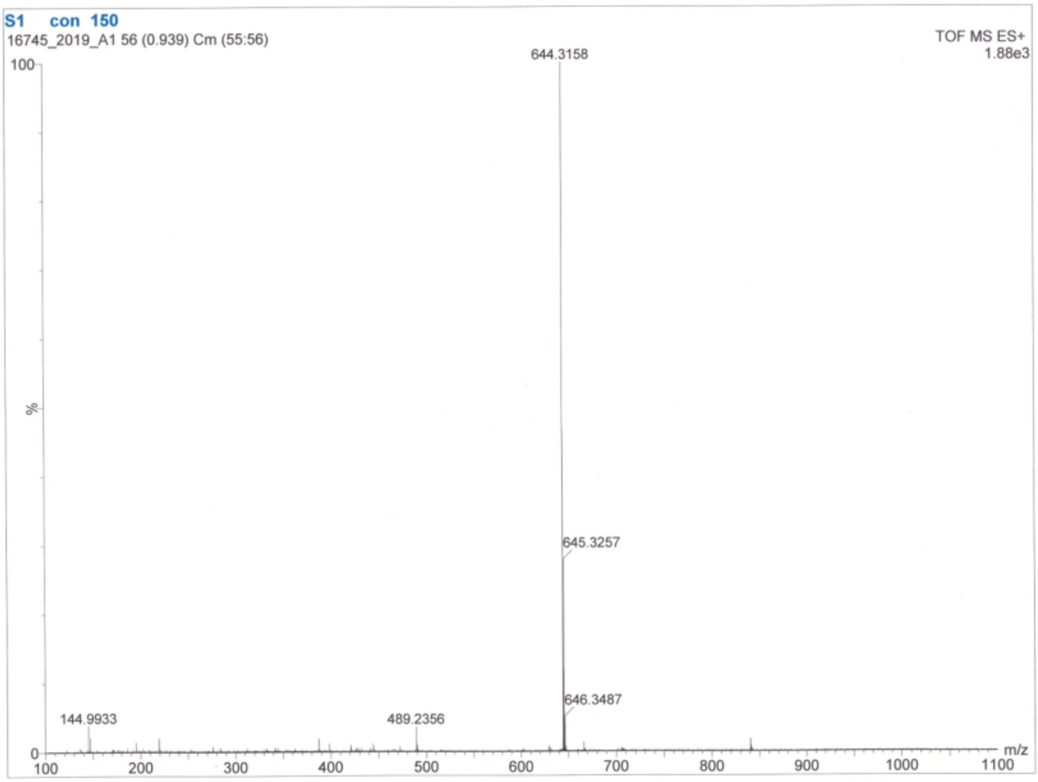

**Ac-Abu(Bth)-Dap-Gly-Gly-ACC**

HRMS for $C_{31}H_{34}N_8O_8S$ ($m/z_{calcd}$ = 679.23; $m/z_{found}$ = 679.12)

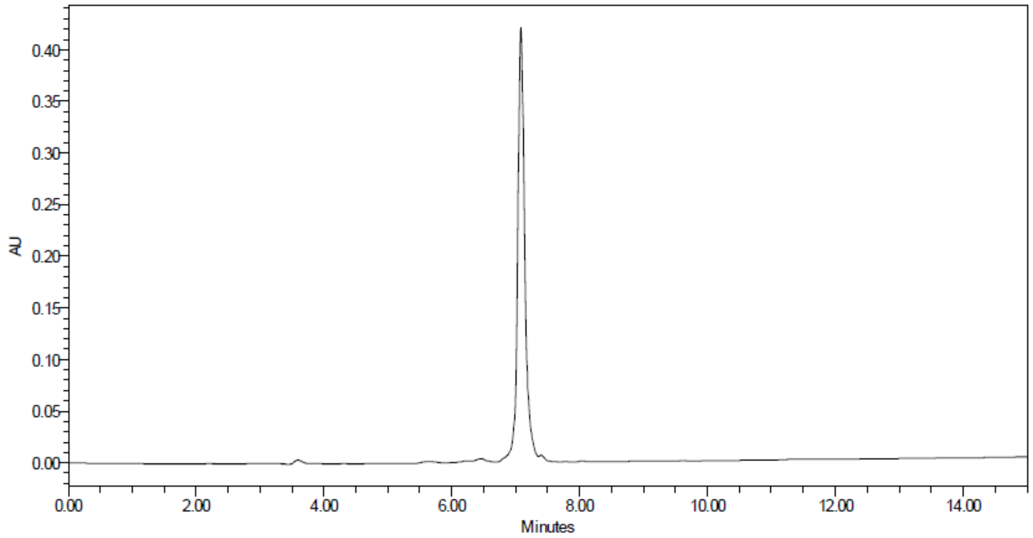

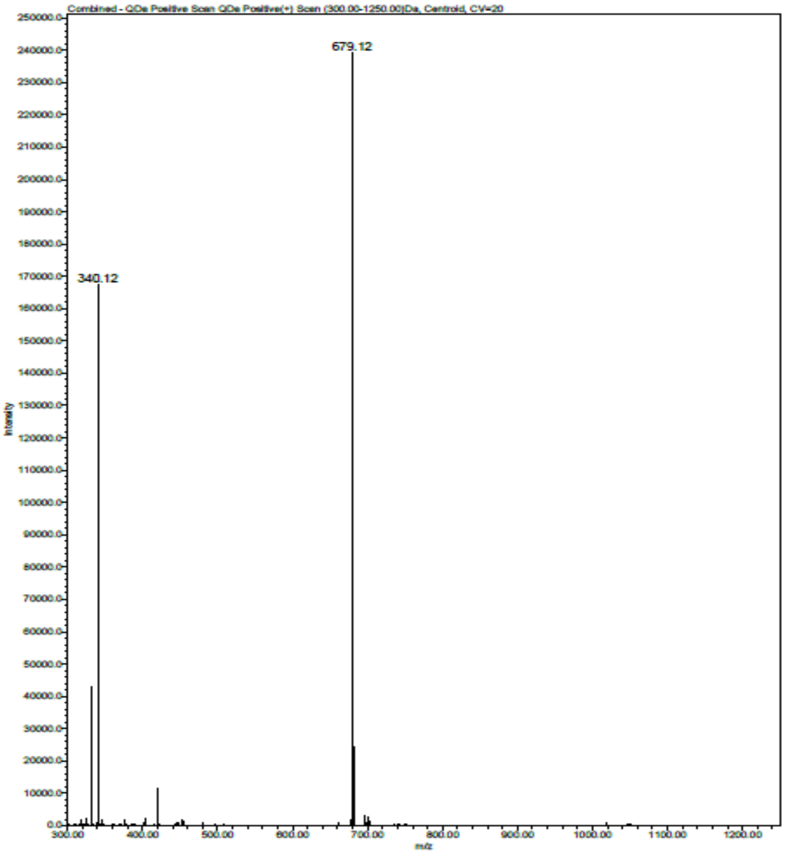

**Ac-Abu(Bth)-Dab-Gly-Gly-ACC**

HRMS for $C_{32}H_{36}N_8O_8S$ (m/$z_{calcd}$ = 693.25; m/$z_{found}$ = 693.14)

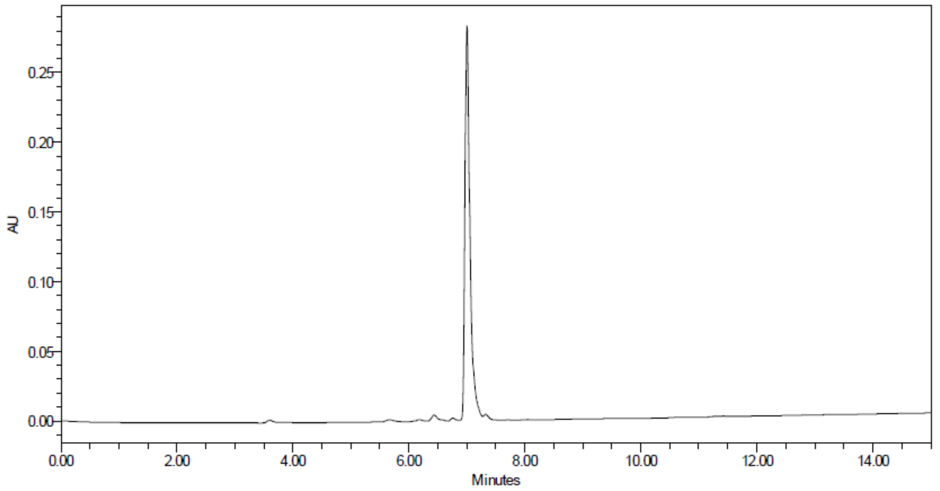

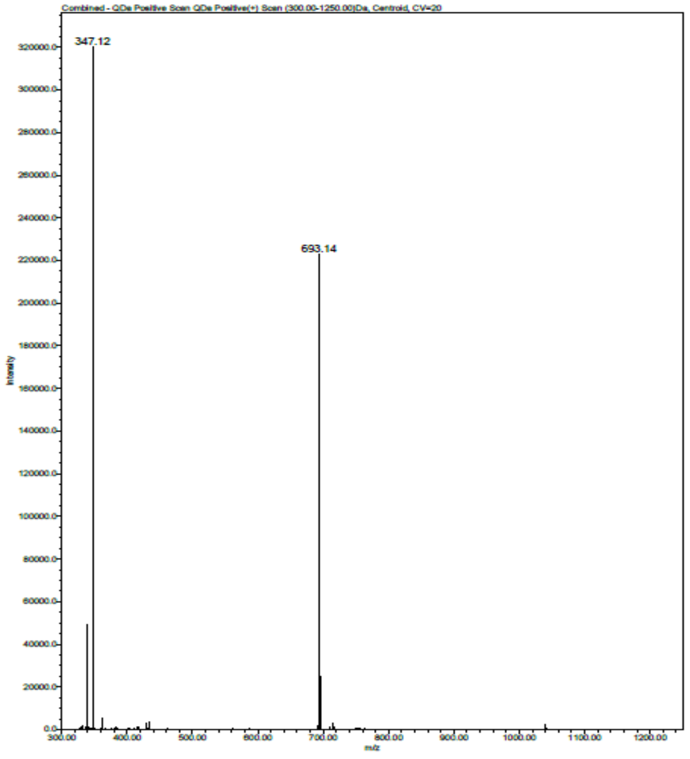

**Ac-hTyr-Phe(guan)-Gly-Gly-ACC**

HRMS for $C_{37}H_{41}N_9O_9$ ($m/z_{calcd}$ = 756.31; $m/z_{found}$ = 756.23)

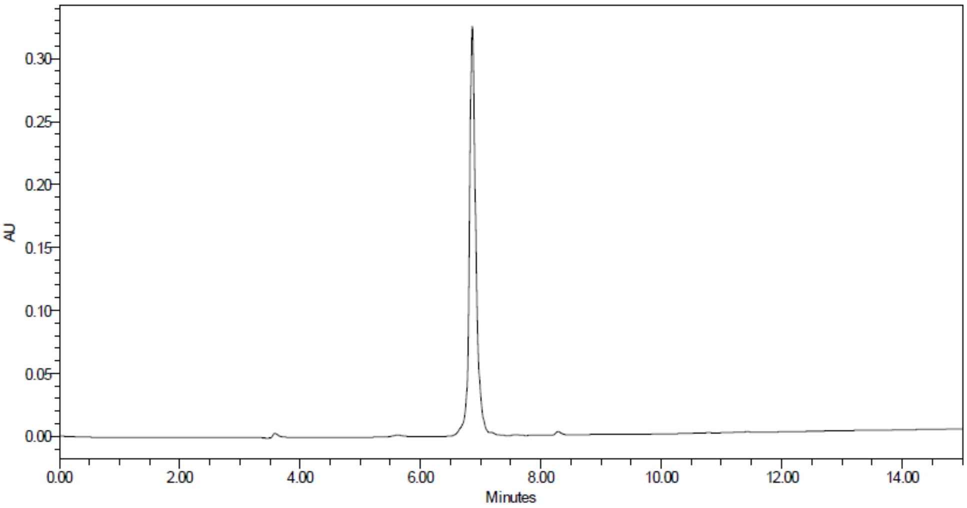

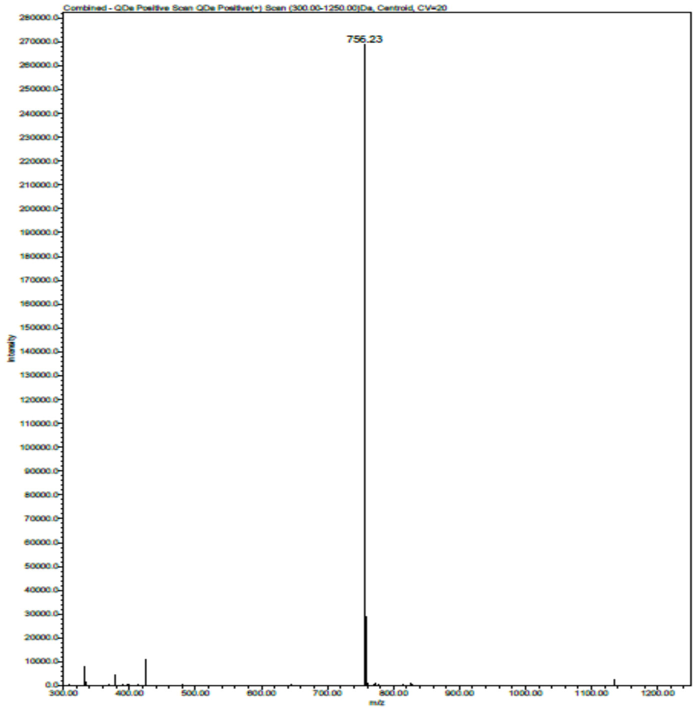

**Ac-hTyr-Dap-Gly-Gly-ACC**

HRMS for $C_{30}H_{35}N_7O_9$ ($m/z_{calcd}$ = 638.26; $m/z_{found}$ = 638.12)

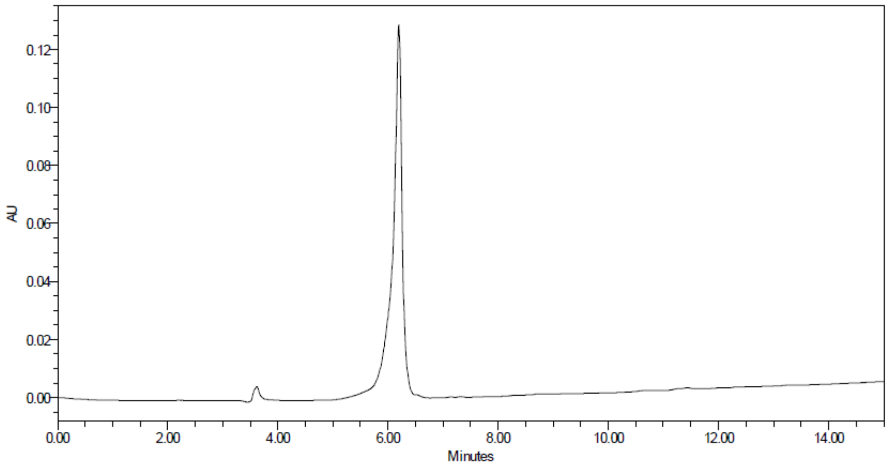

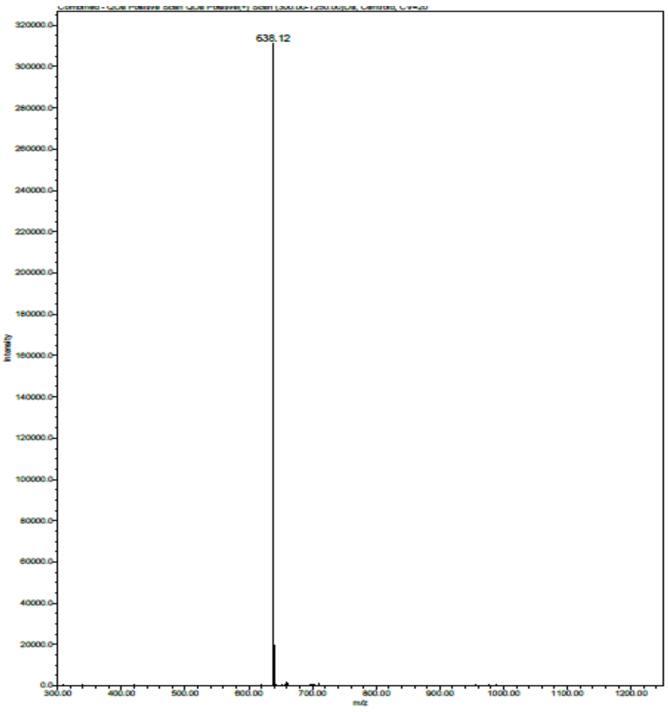

**Ac-hPhe-Dap-Gly-Gly-ACC**

HRMS for $C_{30}H_{35}N_7O_8$ ($m/z_{calcd}$ = 622.26; $m/z_{found}$ = 622.12)

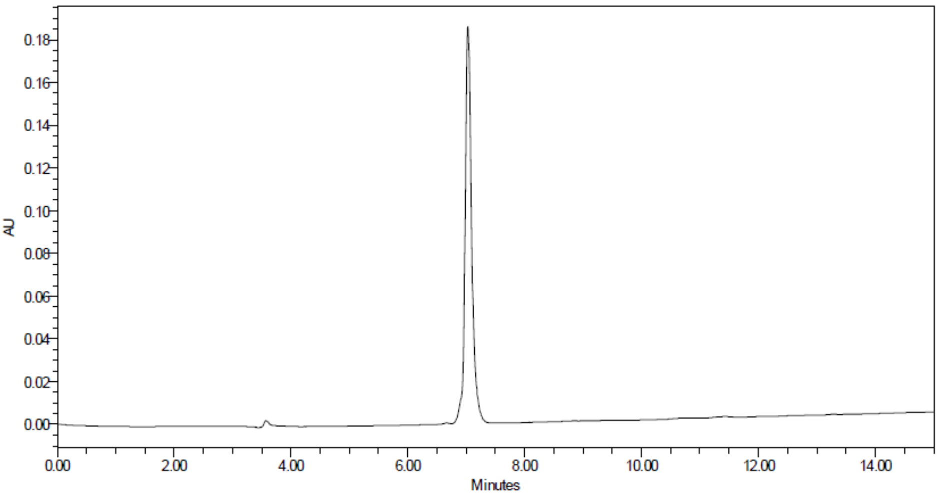

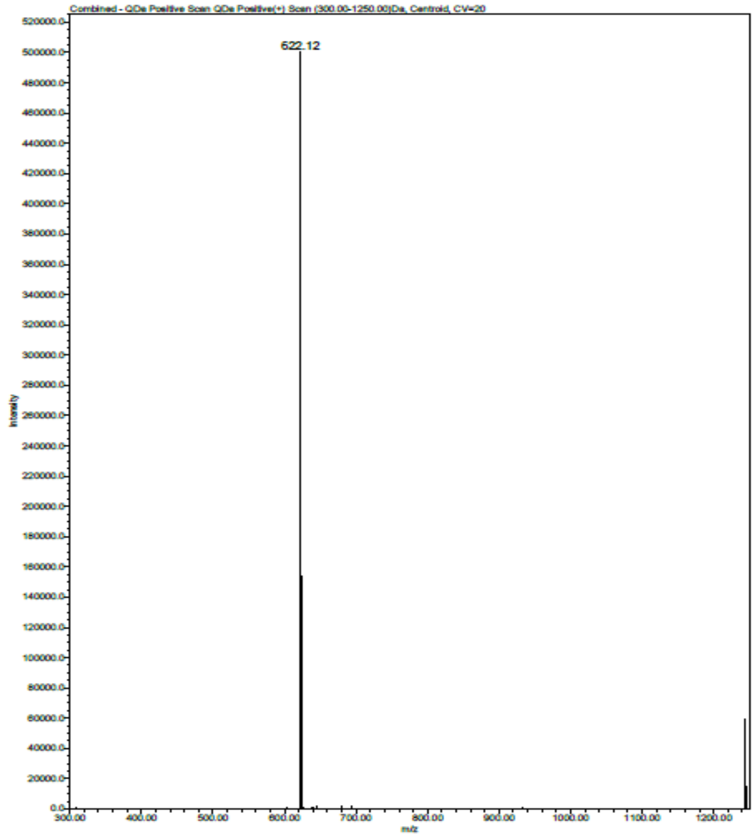

**Ac-Abu(Bth)-Dap-Gly-Gly-VME**

HRMS for $C_{23}H_{30}N_6O_6S$ ($m/z_{calcd}$ = 519.2026; $m/z_{found}$ = 519.2034)

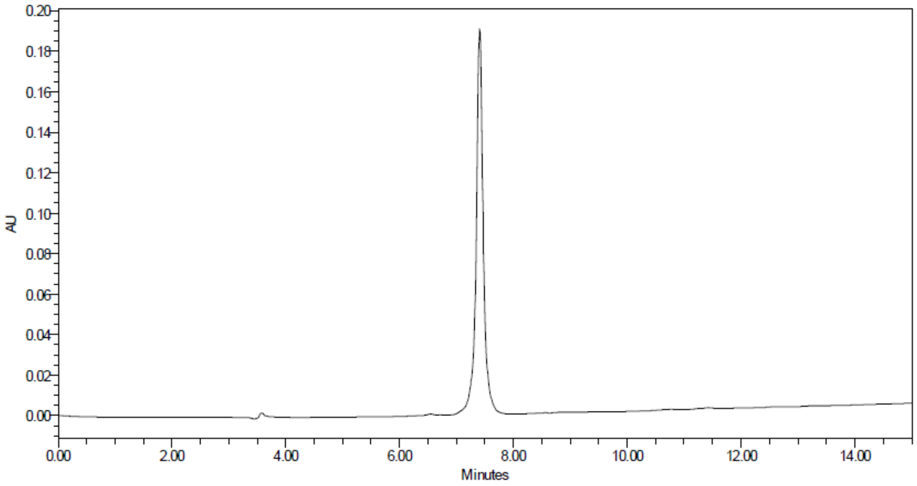

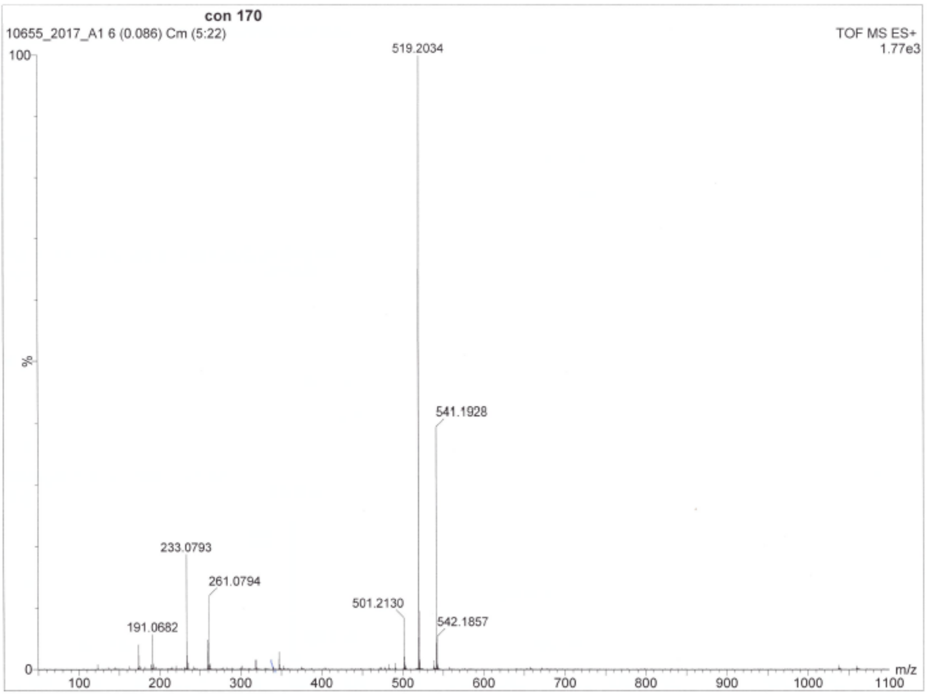

**Ac-hTyr-Dap-Gly-Gly-VME**

HRMS for $C_{22}H_{31}N_5O_7$ ($m/z_{calcd}$ = 478.2302; $m/z_{found}$ = 478.2291)

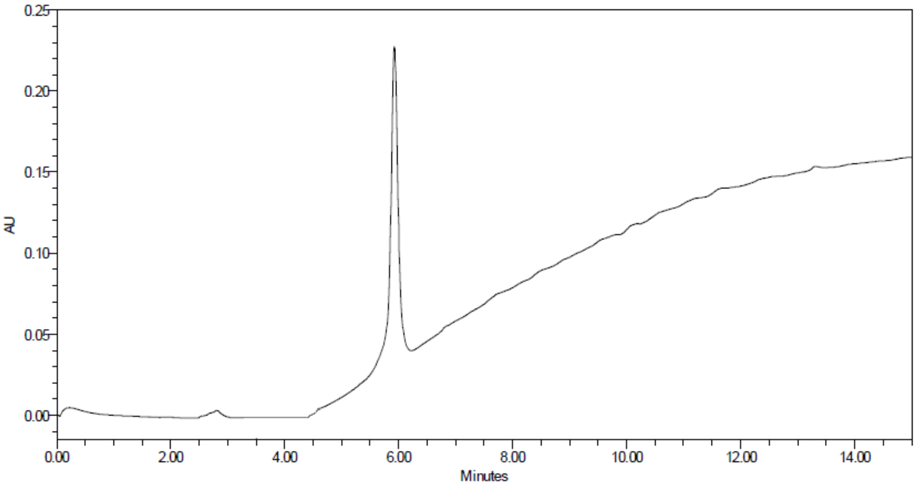

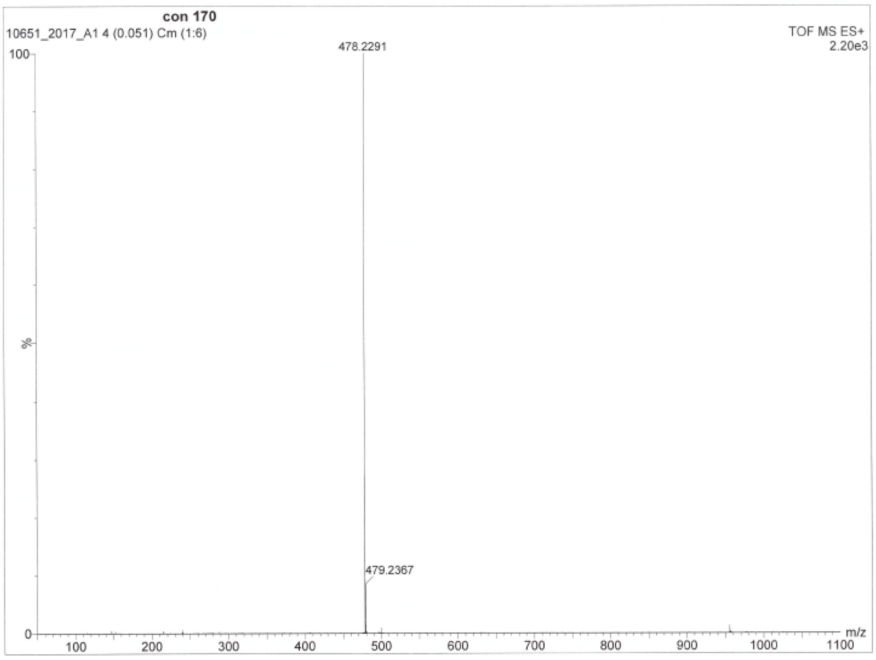

**Ub-ACC**

HRMS for $C_{389}H_{637}N_{107}O_{120}S$

| charge | m/z$_{calcd}$ | m/z$_{found}$ |
|---|---|---|
| +6 | 1461.6221 | 1461.8839 |
| +7 | 1252.9628 | 1253.2618 |
| +8 | 1096.4684 | 1096.7080 |
| +9 | 974.7505 | 974.9593 |
| +10 | 877.3762 | 877.6428 |
| +11 | 797.7063 | 797.8460 |

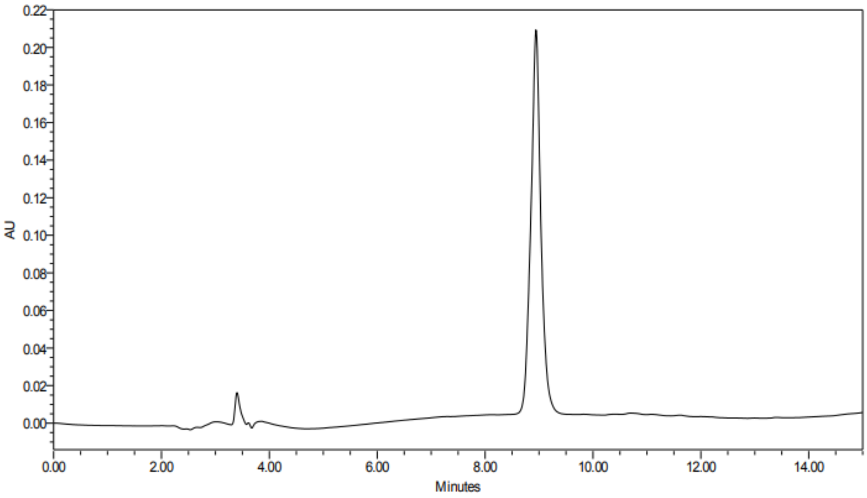

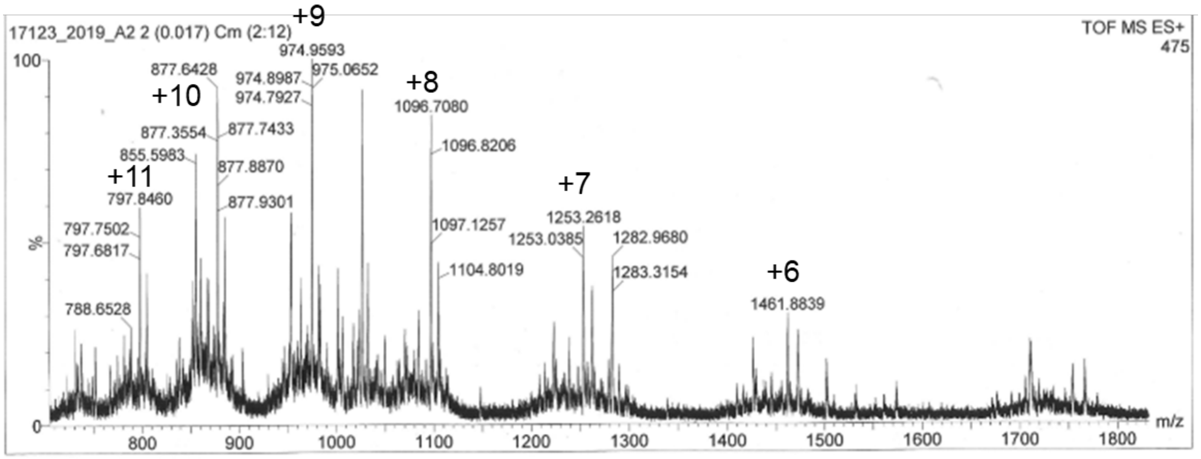

**Biot-6-Ahx-Ub-VME**

HRMS for $C_{397}H_{658}N_{108}O_{121}S_2$

| charge | $m/z_{calcd}$ | $m/z_{found}$ |
|---|---|---|
| +6 | 1491.4778 | 1491.8063 |
| +7 | 1278.5535 | 1278.8268 |
| +8 | 1118.8602 | 1119.2485 |
| +9 | 994.6543 | 994.8820 |
| +10 | 895.2896 | 895.3971 |
| +11 | 813.9912 | 814.0791 |
| +12 | 746.2426 | 746.3508 |

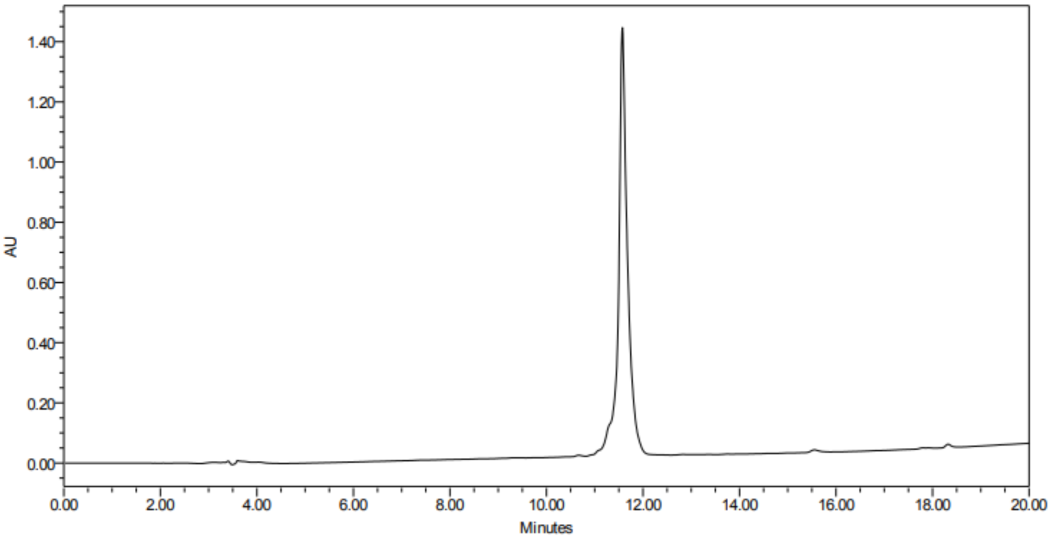

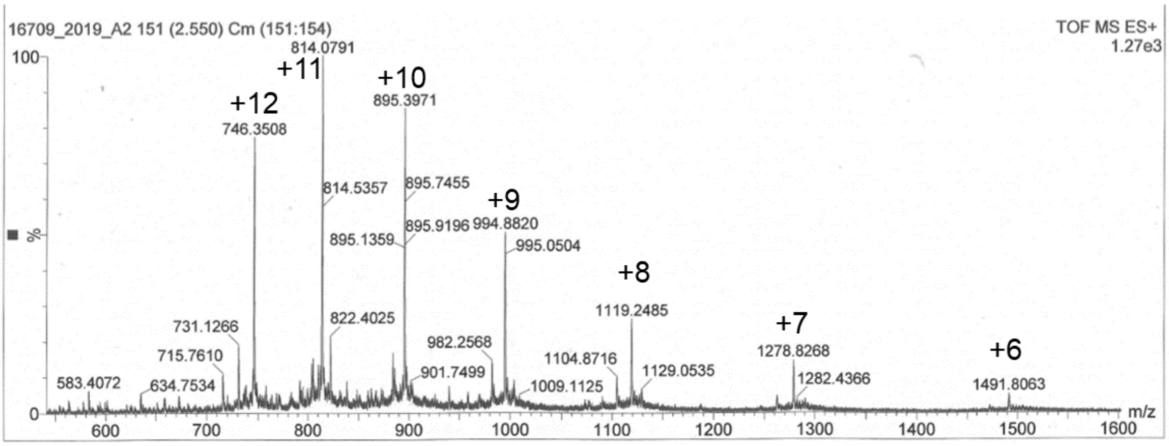
